# High frequency neuronal bursting is essential for circadian and sleep behaviors in *Drosophila*

**DOI:** 10.1101/2020.04.20.051441

**Authors:** Florencia Fernandez-Chiappe, Lia Frenkel, Carina Celeste Colque, Ana Ricciuti, Bryan Hahm, Karina Cerredo, Nara Inés Muraro, María Fernanda Ceriani

**Affiliations:** Instituto de Investigación en Biomedicina de Buenos Aires (IBioBA) - CONICET - Partner Institute of the Max Planck Society, Buenos Aires, Argentina; Fundación Instituto Leloir - IIBBA - CONICET, Buenos Aires, Argentina; Instituto de Biociencias, Biotecnologías y Biomedicina (IB3) - Departamento de Fisiología Biología Molecular y Celular - Facultad de Ciencias Exactas y Naturales - Universidad de Buenos Aires, Buenos Aires, Argentina

**Author notes:** These authors contributed equally to this work (Florencia Fernandez-Chiappe, Lia Frenkel). Shared corresponding authors. Correspondence.

## Abstract

Circadian rhythms have been extensively studied in *Drosophila*, however, still little is known about how the electrical properties of clock neurons are specified. We have performed a behavioral genetic screen through the downregulation of candidate ion channels in the lateral ventral neurons (LNvs) and show that the hyperpolarization-activated cation current I_h_ is important for the behaviors that the LNvs command: temporal organization of locomotor activity and sleep. Using whole-cell patch clamp electrophysiology we demonstrate that small LNvs are bursting neurons, and that I_h_ is necessary to achieve the high frequency bursting firing pattern characteristic of both types of LNvs. Since firing in bursts has been associated to neuropeptide release, we hypothesized that I_h_ would be important for LNvs communication. Indeed, herein we demonstrate that I_h_ is fundamental for the recruitment of PDF filled dense core vesicles to the terminals at the dorsal protocerebrum and for their timed release, and hence for the temporal coordination of circadian behaviors.

## Introduction

Circadian (*circa*: around, *diem:* day) rhythms are biological rhythms with a period of approximately 24h that have evolved in essentially all organisms. They confer an important adaptive value by allowing the anticipation to the daily changes in environmental conditions associated to the rotation of our planet. The “around the clock” coordination of behavior and physiology in *Drosophila* is regulated by approximately 150 neurons grouped in different clusters and named after their anatomical localization (1). Among them, the small lateral ventral neurons (sLNvs) have been identified as a fundamental group in the control of behavioral rhythms under free running conditions, communicating via the release of the neuropeptide Pigment Dispersing Factor (PDF) (2-5) and glycine (6). The large lateral ventral neurons (lLNvs), on the other hand, are highly relevant for arousal and the PDF they release provides wake promoting functions (7-9).

Although the mechanisms that give rise to the cell-autonomous cycling of gene and protein expression and comprise the core of the molecular clock have been described thoroughly (10), one of the challenges of the field now is to understand how different clock neurons communicate to each other. It is indeed the emerging properties of these clock neuronal circuits acting concertedly that provide the system with plasticity and adaptability (11). But to understand the communication taking place within clock neurons, it is paramount to examine the physiological properties of the different neuronal groups. The type, amount and distribution of ion channels present in the membrane of a neuron determine features such as excitability and action potential firing pattern. In particular, clock neurons change their electrical activity on a daily basis, with higher action potential firing during the day than at night, a phenomenon that has been described both in mammals and flies (reviewed in (12).

In *Drosophila*, several ion channels have already been found to play roles in different aspects of circadian function, such as the calcium dependent voltage-gated potassium channel *slowpoke* (*slo*) (13, 14) and its binding protein (*slob*) (13, 15, 16), the cation channel *narrow abdomen* (*na*) (17-19), the voltage-gated potassium channel *Shaw* (20, 21), the inward rectifying potassium channel *Ir* (22), the temperature sensitive *trpA1* channel (23), the potassium channel *hyperkinetic* (*hk*) (24) and the voltage-gated potassium channel *Shal* (21, 25). Under the hypothesis that additional ion channels are involved in determining the characteristic physiological properties of the LNvs that ensure circadian organization of locomotor activity, we performed a behavioral genetic screen downregulating candidate ion channels using RNA interference (RNAi) specifically in LNvs. Following this strategy, we have been able to identify several ion channels that, when knocked down, alter circadian locomotor behavior under free running conditions. Of those, we have first focused our attention on the hyperpolarization-activated cation current I_h_, since, as it has been described in mammalian neurons (26), its biophysical properties make it particularly suitable to mediate the organization of action potential firing in bursts, a firing mode that characterizes lLNvs (27-29) and, we show here, also sLNvs. Consistently, we demonstrate that perturbing *I_h_* causes a decrease in the frequency of LNvs bursting that is accompanied by a reduction in PDF immunoreactivity and in the complexity of sLNv axonal termini. Moreover, we have found that the disruption of *I_h_* is accompanied by an increase in sleep. Altogether, our results reveal a novel function of I_h_ in determining LNvs physiology and the behaviors they command, and uncover several additional ion channels with putative roles in these important clock clusters, for future exploration.

## Results

### LNvs ion channel downregulation behavioral screen

To shed light onto how LNvs achieve the physiological properties that allow them to play a key role in the circadian organization of locomotor activity, we performed an ion channel downregulation behavioral screen. The *pdf-Gal4* driver, in the presence of *UAS-dicer2* (from here on *pdf,dicer*) was used to drive expression of UAS-RNAis to knock down the expression of candidate ion conductances solely in LNvs. The RNAis were chosen to target ion channel genes, genes coding for ion channel auxiliary subunits or genes coding for ion channel transporters which had not been reported to be involved in LNvs-driven circadian phenotypes before. The locomotor activity of *pdf,dicer>RNAi* male flies was recorded using *Drosophila* Activity Monitors (DAM, Trikinetics) for 9 days in DD after 3 days of LD entrainment. Each RNAi was initially tested once and, in the case of showing a differential phenotype in DD, corresponding to either a change of circadian period or deconsolidation of locomotor activity, experiments were repeated. In some cases a non-significant trend towards a phenotype was detected and therefore two RNAis that targeted different regions of the same gene were genetically combined to achieve a more potent downregulation. **Table 1** shows the positive hits of our screen, revealing novel ion channels or ion channel auxiliary subunits likely to play roles in LNvs circadian function, namely: *cacophony* (*cac*, CG1522), *Ca-α1T* (*Ca^2+^-channel protein α_1_ subunit T*, CG15899), *ClC-a* (*Chloride channel-a*, CG31116), *CngA* (*Cyclic nucleotide-gated ion channel subunit A*, CG42701), *I_h_* (*I_h_ channel*, CG8585), *Ork1* (*Open rectifier K^+^ channel* 1, CG1615), *Shal* (*Shaker cognate l*, CG9262) and *tipE* (*temperature-induced paralytic E*, CG1232). The RNAis that did not show altered circadian phenotypes in our screen are listed in **S1 Table**. In total 70 RNAis aimed at 36 different genes were tested.

**Table 1:**
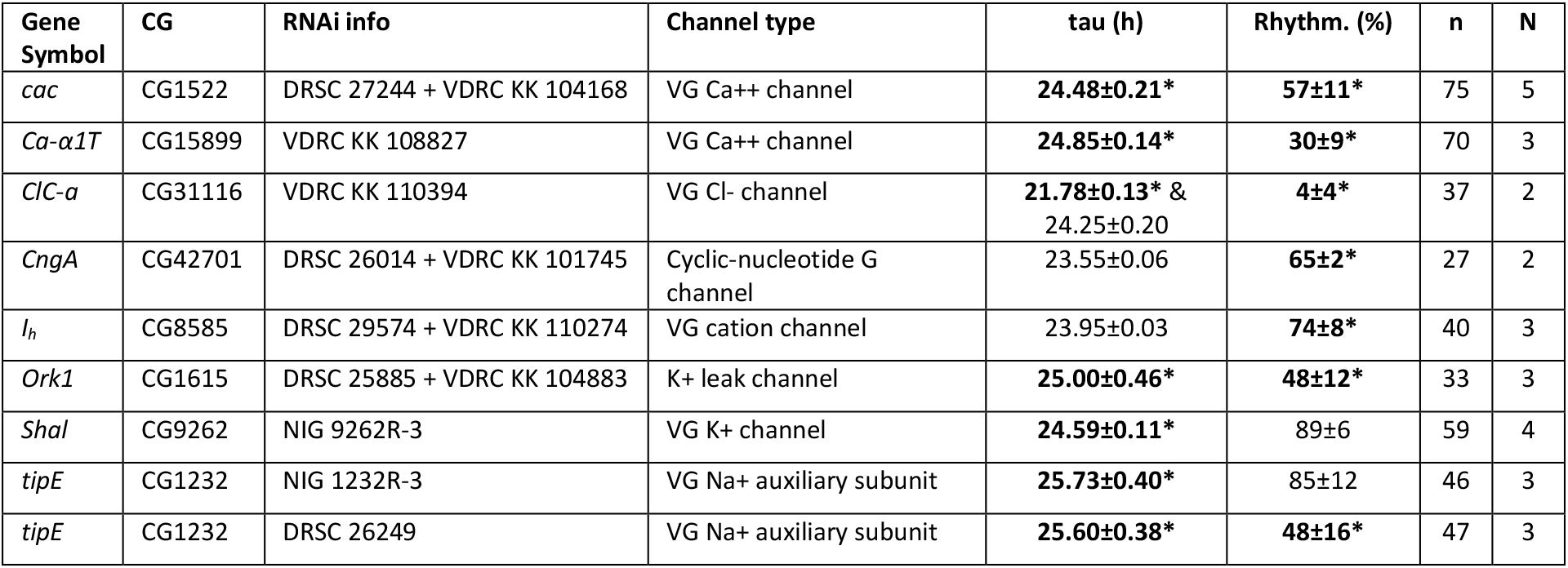
Positive hits of the ion channel downregulation behavioral screen. This table includes a list of genes that, when downregulated exclusively in LNvs using these particular RNAi constructs, produced statistically significant alterations in free running period and/or percentage of rhythmicity. Values represent the average of mean values of N independent experiments ± SEM. n indicates total number of individuals tested. * Indicates statistically significant difference (p<0.05) after a one-way ANOVA comparing *pdf,dicer>RNAi* to control genotypes *pdf,dicer/+* (average period 24.00±0.07h and average rhythmicity 95±3%) and RNAi/+ (average period 24.08±0.17h and average rhythmicity 93±4%). Tukey test was used for means comparison and Levene’s test for checking ANOVA assumption of homogeneity of variance. In the case where information for two RNAi constructs is given, it means that each RNAi on its own did not show significant differences compared to controls, but did show a trend towards an altered phenotype. For that reason two different RNAis for the same gene were genetically combined to achieve added downregulation strength. In the case of *ClC-a*, the reduction of rhythmicity was due to the appearance of complex rhythms and not to the deconsolidation of locomotor activity organization; the tau of each component of complex rhythms is given. V: Voltage, G: Gated, DRSC: *Drosophila* RNAi Screening Center, VDRC: Vienna *Drosophila* Resource Center, NIG: National Institute of Genetics.

Although all of the ion channel genes that resulted positive hits of our behavioral screen are worth of further assessment, we focused our attention on the hyperpolarization-activated cation current I_h_. Little is known about this channel in *Drosophila*, but its homologues in mammals have been implicated in diverse functions such as the generation of pacemaker potentials and the determination of neuronal excitability, among others (26). RNAi-mediated downregulation of *I_h_* in LNvs produced a subtle but consistent decrease in locomotor rhythmicity without altering free running period (**Tables 1 and 2, Fig 1A**).

**Table 2:** *I_h_* genetic manipulations disrupt circadian locomotor activity organization. DD Analysis (left): the average free running period and percentage of rhythmicity of populations of male flies of the indicated genotypes are shown. Values represent the average of N independent experiments ± SEM. n indicates total number of individuals tested. *Indicates statistically significant difference (p<0.05) after a one-way ANOVA comparing experimental genotypes to genetic controls. 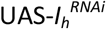 refers to the genetic combination of two 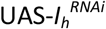 constructs: DRSC 29574 + VDRC KK 110274 mentioned in Table 1). In the case of the *I_h_* null mutants, 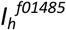 and 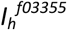, homozygotes were compared to a *w^1118^* control and to heterozygotes (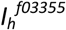 crossed by *w^1118^*). RU refers to the presence of the steroid RU486 (200 μg/ml), the activator of the GeneSwitch system, in the food media. LD Analysis (right): Morning Anticipation Index (MAI) and Evening Anticipation Index (EAI) were calculated for the same genotypes. *Indicates statistically significant difference (p<0.05) after Kruskal-Wallis statistical analysis with Dunn’s multiple comparisons test.

| Genotype | DD Analysis |  |  |  | LD Analysis |  |  |  |
| --- | --- | --- | --- | --- | --- | --- | --- | --- |
|  | tau (h) | Rhythm.(%) | n | N | MAI | EAI | n | N |
| *pdf,dicer/+* | 24.00±0.07 | 95±3 | 75 | 3 | 0.73±0.02 | 0.88± 0.02 | 55 | 2 |
| *UAS-I<sub>h</sub><sup>RNAi</sup>/+* | 23.71±0.02 | 92±6 | 39 |  | 0.77±0.03 | 0.80±0.03 | 21 |  |
| *pdf,dicer>UAS-I<sub>h</sub><sup>RNAi</sup>* | 23.95±0.03 | **74±8\*** | 40 |  | 0.67±0.03 | 0.80±0.02 | 23 |  |
| *pdfGS/+ , RU* | 24.59±0.41 | 97±2 | 64 | 3 | 0.64±0.01 | **0.66±0.02\*** | 76 | 3 |
| *UAS-I<sub>h</sub><sup>RNAi</sup>/+ , RU* | 23.75±0.06 | 100±0 | 41 |  | 0.72±0.02 | 0.58±0.01 | 50 |  |
| *pdfGS>UAS-I<sub>h</sub><sup>RNAi</sup> , RU* | 24.81±0.75 | **57±12\*** | 67 |  | **0.59±0.01\*** | 0.60±0.01 | 74 |  |
| *control* | 24.02±0.05 | 96±2 | 86 | 4 | 0.68±0.03 | 0.84±0.02 | 62 | 2 |
| *I<sub>h</sub><sup>f01485</sup>/+* | 23.87±0.03 | 96±3 | 80 |  | 0.71±0.04 | 0.89±0.02 | 32 |  |
| *I<sub>h</sub><sup>f01485</sup>* | 23.56±0.05 | **60±3\*** | 98 |  | 0.55±0.05 | **0.65±0.03\*** | 34 |  |
| *I<sub>h</sub><sup>f03355</sup>/+* | 23.86±0.05 | 99±1 | 96 |  | 0.73±0.03 | 0.92± 0.02 | 30 |  |
| *I<sub>h</sub><sup>f03355</sup>* | 23.88±0.08 | **39±12\*** | 104 |  | **0.51±0.03\*** | **0.67±0.03\*** | 30 |  |

**Fig 1:**
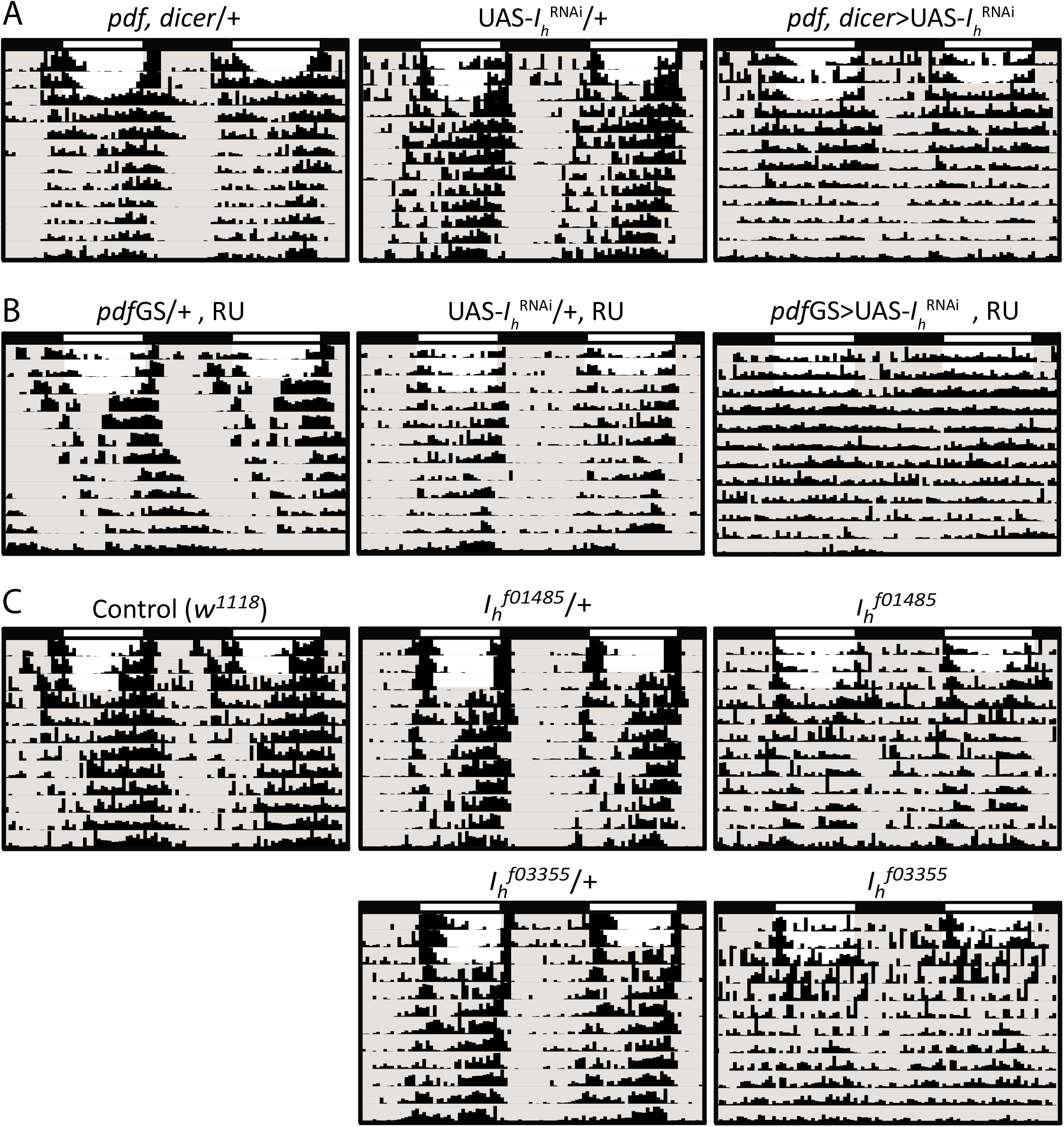
*I_h_* genetic manipulations disrupt circadian locomotor activity organization. Representative double-plotted actograms of the different *I_h_* genetic manipulations tested. **A)** LNvs constitutive downregulation of *I_h_* using *pdf,dicer* and 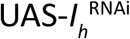 (in all cases 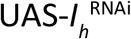 refers to the genetic combination of two 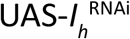 constructs: DRSC 29574 + VDRC KK 110274 mentioned in Table 1) and genetic controls. **B)** LNvs acute downregulation of *I_h_* using pdfGS and 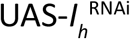 and genetic controls. RU refers to the presence of the steroid RU486, the activator of the GeneSwitch system, in the food media. **C)** Homozygote *I_h_* null mutants, 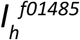 and 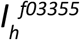, and controls *(w^1118^* and heterozygote mutants, crossed by *w^1118^*). In the case of the experimental genotypes an actogram of an arrhythmic individual is shown, different genetic manipulations varied in the degree of arrhythmicity (see Table 2). No statistically significant alterations in free running period were found for these genetic manipulations.

The *pdf*-Gal4 driver used for the ion channel behavioral screen is active throughout development. Therefore, to dissect whether the behavioral phenotype observed was due to a developmental defect or to a post-developmental functional role, we downregulated *I_h_* expression in LNvs in an adult-specific fashion using the GeneSwitch inducible system (30). When the previously reported *pdf*-GeneSwitch (*pdf*GS) driver (31) was utilized to knock down *I_h_* adult-specifically in LNvs, we also observed a decrease in circadian rhythmicity (**Table 2, Fig 1B**), indicating that the I_h_ channel is necessary post-developmentally in LNvs for the maintenance of circadian function. As a complementary approach, we assessed the circadian behavior of previously reported *I*h null mutants, *I_h_^f01485^* and *I_h_^f03355^* (32, 33). As expected, these mutants also showed reduced rhythmicity under free running conditions (**Table 2, Fig 1C**). Not surprisingly, *I_h_* mutants are less rhythmic than any tissue-specific knock down (LNvs-specific manipulations), strongly suggesting that I_h_ may be necessary not only in LNvs but also in other neuronal types for the rhythmic organization of locomotor activity under free running conditions. All these genetic manipulations did not, in any case, produce changes in free running period (**Table 2**). To assess if I_h_ is important for circadian function also under entrained conditions, we analyzed morning and evening anticipatory behavior. Consistent with the strength of the phenotypes observed in DD, we detected a failure in both, morning and evening anticipation in 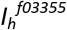 mutants, which is less pronounced in the 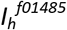 mutants and the adult-specific downregulation of *I_h_*; no effects were detected under *I_h_* constitutive knock down, suggesting potential compensatory mechanisms (**Table 2**). Taking together, these results suggest that the *I_h_* channel contributes to define the firing properties of neurons controlling circadian behavior.

### *I_h_* is necessary for high frequency bursting firing of LNvs

One of the main reasons why we decided to select *I_h_* as the ion channel for in depth analysis is the association of *I_h_* with the organization of action potential firing in bursts. It has been reported, mainly from mammalian thalamic relay and inferior olivary nucleus neurons, that a combination of a hyperpolarization-activated cation current such as I_h_, together with a low-voltage activated T-type calcium current (a channel type also uncovered by our screen, **Table 1**), could mediate a bursting firing mode (26). This is because of I_h_ particular biophysical properties, which opens upon hyperpolarization but carries a depolarizing current (mainly due to the influx of Na^+^). This current takes the membrane potential to the activation threshold of the T-type voltage-gated Ca^++^ channel which depolarizes the membrane up to the action potential firing threshold, opening the classical voltage-gated Na^+^ channels. Because I_h_ is slow to close and does not inactivate, the membrane stays in a depolarized state for longer, generating a burst of action potentials. Once I_h_ closes, the classical voltage-gated K^+^ channels that repolarize the membrane, together with the leak K^+^ channels, produce the after-hyperpolarization that kick starts the following burst, activating I_h_ again (26).

Another relevant observation is that firing in bursts is an effective way of releasing neuropeptides, which are stored in dense core vesicles. In contrast to small clear vesicles containing classical fast neurotransmitters, neuropeptide-filled dense core vesicles require a larger amount of Ca^++^ entering the cell to activate the machinery that fuses and releases the dense core vesicles content (reviewed in (34, 35)). Since lLNvs have been described to fire action potentials in a bursting mode (27, 28) and also to be neuropeptide-releasing neurons (2, 36), the hypothesis we formulated is that I_h_ participates in the active bursting firing mode of LNvs and plays a role in the release of PDF.

We first tested our hypothesis in the lLNvs, which have effectively been shown to be bursting neurons (27, 28). We performed *ex vivo* whole-cell current clamp recordings of control *pdf*-RFP (expressing a red fluorofore in the LNvs thus enabling the identification of the two neuronal types due to the difference in the size of their soma) lLNvs and compared their bursting frequency to 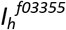 homozygote mutants. Because we have reported that lLNv bursting frequency also depends on synaptic inputs that are disrupted during the dissection protocol (29) we compared the bursting frequency of control and 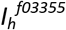 mutant lLNvs at exactly the same time post-dissection (23 minutes). **Fig 2A and D** show that although lLNvs from 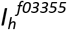 homozygote mutants can still organize their action potential firing in bursts, they do so at a statistically significant lower frequency (mean bursting frequency ± SEM (bursts/min) are lLNvs_CONTROL_=26.2±0.9 and lLNvs_*Ihf03355*_=17.5±1.4). Other parameters, such as overall firing frequency and membrane potential were not significantly affected in 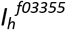 mutants (**S1 Fig A and B**).

**Fig 2:**
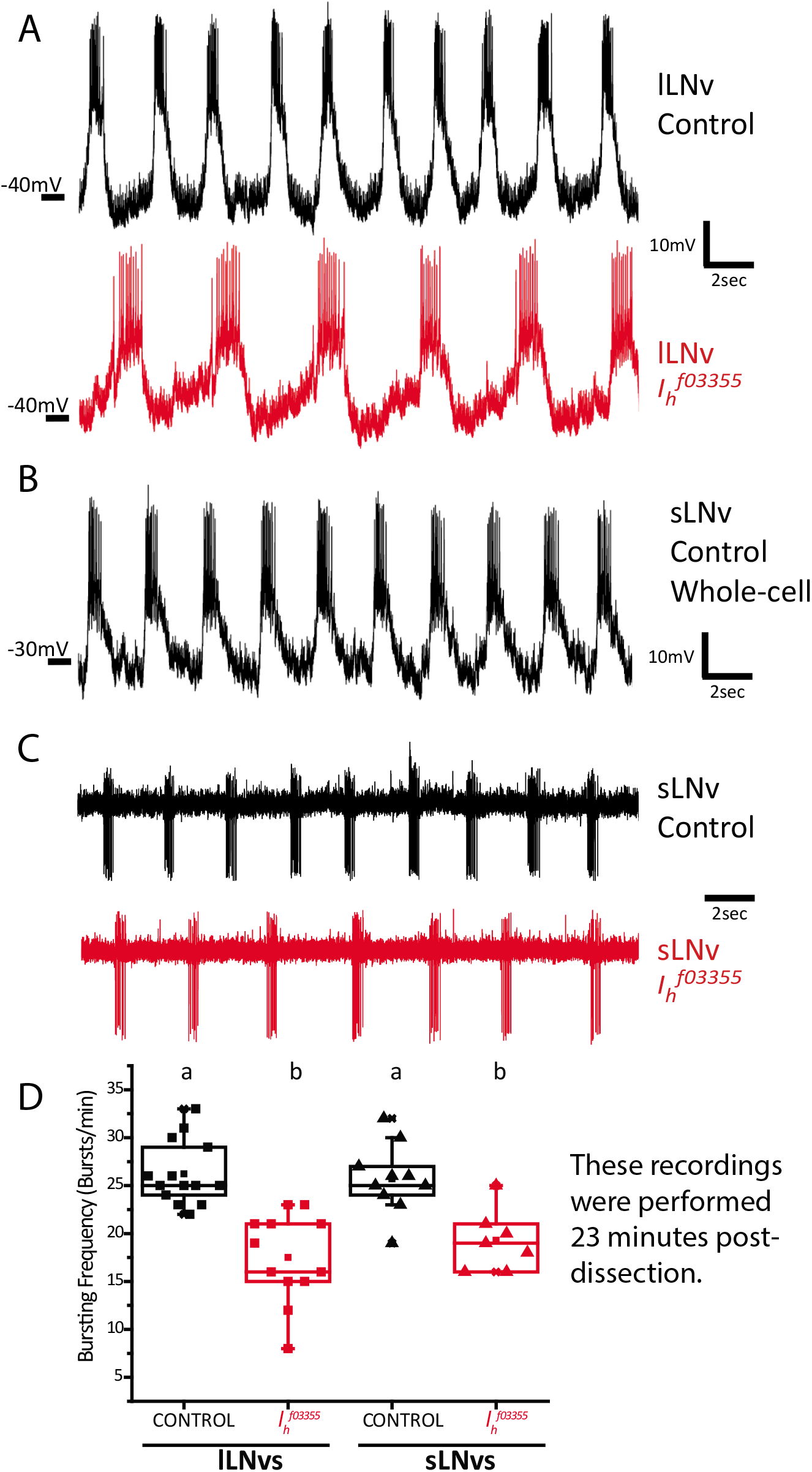
*I_h_* is important for high frequency bursting of LNvs. **A)** Representative traces of whole-cell patch clamp recordings of lLNvs of control *(pdf-RFP*, top) and *I_h_* homozygote mutant genotypes (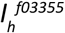; *pdf*-RFP, bottom). **B)** Representative trace of a recording of a sLNv control (*pdf*-RFP) in whole-cell patch clamp configuraton. **C)** Representative traces of cell-attached recordings of sLNvs of control (pdf-RFP, top) and *I_h_* homozygote mutant genotypes (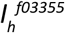; *pdf*-RFP, bottom). **D)** Box plot showing burstng frequency quantficaton of lLNvs and sLNvs of control (*pdf*-RFP) and *I_h_* homozygote mutant genotypes (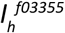; *pdf*-RFP). All quantficatons were done at exactly 23min post-dissecton. Different letters indicate significant differences (p<0.05) after a one-way ANOVA with Tukey test for means comparisons. n: lLNvs_CONTROL_=14, lLNvs_Ihf03355_=12 SLNvs^CONTROL^=10 SLNvs_Ihf03355_=7.

Next, we tested our hypothesis in the sLNvs. Information regarding sLNvs electrophysiological properties is scarce (28, 37), probably due to the technical challenge that their small soma size represents. However, given the important role that sLNvs play in the control of circadian behavior, we analyzed their firing properties in detail. We report here that the sLNvs also fire action potentials organized in bursts (**Fig 2B**). Because obtaining a large amount of recordings in whole-cell configuration was a difficult task to achieve, we recorded action potential firing rate and bursting frequency in a cell-attached configuration of the sLNvs of control (*pdf*-RFP) and 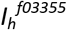 homozygote mutants. We found that, similarly to lLNVs, sLNvs show a decreased bursting frequency in the absence of *I_h_* (**Fig 2C and D,** mean bursting frequency ± SEM (bursts/min) are sLNvs_CONTROL_=25.7±1.2 and sLNvs*Ihf03355*=19.3±1.3), without significantly affecting overall firing frequency (**S1 Fig C**).

A feature that should be remarked is that both types of LNvs display equivalent basal bursting frequencies (**Fig 2D**), suggesting that this parameter depends on common synaptic inputs and/or shared intrinsic mechanisms. We have previously reported that lLNvs bursting frequency relies to some extent on synaptic inputs coming from the visual neuropiles, which indirectly involve L2 lamina neurons and the neurotransmitter acetylcholine (29). The dependence of lLNv bursting on these synaptic inputs is illustrated by the fact that this parameter decays as a function of the time elapsed since brain dissection, which removes the retina (29). We found that sLNv bursting frequency also decays with time *ex vivo*, which can be seen both at the population level (**S2 Fig A**) and also in individual cells (**S2 Fig B-C**), suggesting that both types of LNvs depend on synaptic inputs which are gradually lost after dissection. Alternatively, it might be that lLNvs rely on visual circuit inputs to burst, and sLNv bursting depends on lLNv bursting. Certainly, the neuronal processes of lLNvs are better localized, spanning all over the optic lobes, to integrate visual information. However, the sLNvs have been shown to receive direct input from the Hofbauer-Buchner (HB) eyelet extraretinal organ (38), whose integrity may also be compromised during dissection. Whether sLNvs and lLNvs rely on different or similar synaptic inputs to support bursting frequency, or if one LNv group depends on the other to detect synaptic information from visual organs, will require further investigation.

We also compared bursting frequency at a different time post-dissection; interestingly, this analysis showed that both large and small LNvs present equivalent bursting frequency, and the lack of *I_h_* produces a significant reduction of this parameter, which is of the same magnitude in the two LNv groups (**S2 Fig D**, compare to Fig 2D, mean bursting frequency ± SEM (bursts/min) are as following ILNvs_CONTROL_=21.0±0.9, lLNvs_*Ihf03355*_=13.8±0.9, sLNvs_CONTROL_=21.1±1.5, sLNvs_*Ihf03355*_=14.6±0.9). Altogether, our results suggest that both LNv clusters share common mechanisms to control their bursting firing frequency, which appear to be controlled intrinsically, likely involving the I_h_ current, as well as rely on synaptic inputs.

### I_h_ channel and the sLNvs outputs

Over the years it has been demonstrated that communication from the sLNvs to other clock clusters is crucial for coherent circadian behavior under free running conditions (2, 3, 5, 6, 14, 39-41). The rhythmic accumulation of PDF neuropeptide in sLNvs axonal termini has been implicated in this communication, with high immunoreactivity detected in the early morning and low immunoreactivity at night (42). We hypothesized that release of dense core vesicles containing PDF would be affected by the decrease in bursting activity that accompanies *I_h_* downregulation. To test this, we performed anti-PDF immunofluorescence in whole brains of flies with adult-specific downregulation of *I_h_*. **Fig 3A-B** shows that PDF immunoreactivity in controls (*pdfGS/+*) displays the normal cycling pattern; however, upon downregulation of 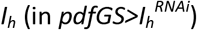 PDF levels at the axonal termini are constantly reduced and clamped in a night-like state.

**Fig 3:**
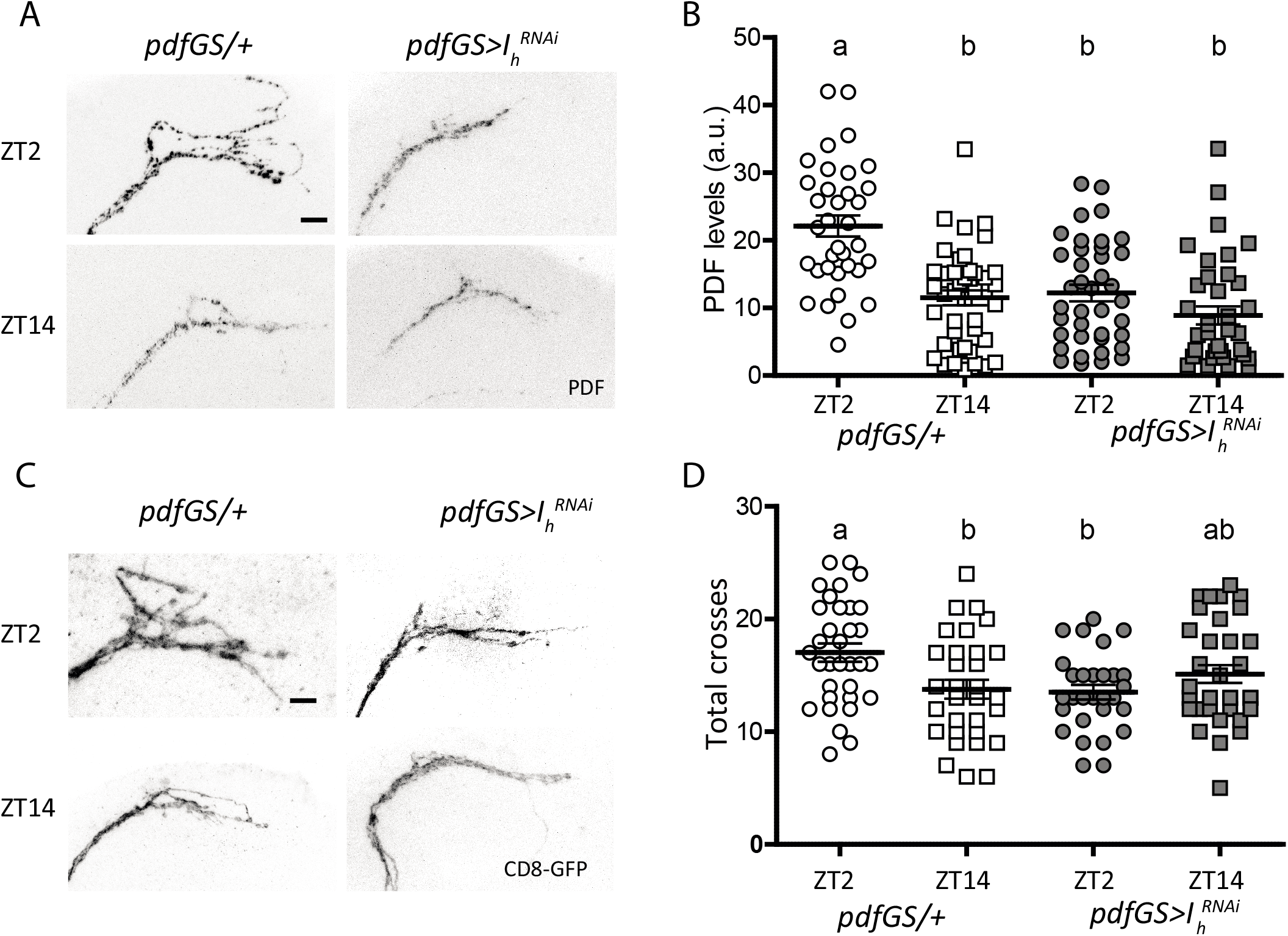
*I_h_* downregulation affects PDF levels and structural plasticity. **A)** Confocal images of representative sLNvs dorsal projections of individual flies of control (*pdf*GS/+) and *I_h_* downregulation 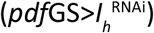 at day (top) and night (bottom) showing their PDF content. Flies were kept in LD 12:12 at 25°C for 7 days in food containing RU486. Brains were dissected at ZT02 and ZT14 and standard anti-PDF immunofluorescence detection was performed. The bar indicates 10μm. **B)** PDF quantitation of the sLNvs dorsal projections for the four conditions mentioned before. Circles represent day time, squares, night time; empty symbols are the control genotype (*pdf*GS/+) and filled symbols, the experimental one 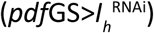. Different letters indicate significant differences, analysis included a two-way ANOVA (genotype and time of day) [F_(3,150)_=18.58 p<0.0001 with Tukey post hoc test, α =0.05], n=35-43 per group. **C)** Confocal images of sLNvs projections illustrating their complexity at ZT02 and ZT14 for both in the control and *I_h_* downregulated genotypes. Procedure as in A but with immunofluorescence against GFP. The bar indicates 10μm. **D)** Complexity quantitation was asses by Sholl analysis (ImageJ) corroborated by visual inspection of each picture. Symbols as in B, analysis included a two-way ANOVA [F_(3,123)_=4.24 p<0.01 with Tukey post hoc test, α =0.05]. Different letters indicate significant differences. n=34-38 per group.

In addition to PDF cyclic accumulation, sLNvs show circadian variation of the complexity of their axonal arborizations (43) in order to contact different synaptic targets at different times of the day (44). This structural synaptic plasticity has been shown to be activity-dependent (31, 45, 46), therefore we wondered whether *I_h_* downregulation would affect this property. **Fig 3C-D** shows that total axonal crosses measured by Sholl analysis in controls display the normal cycling pattern, where the terminals are maximally spread (and more complex) in the early morning and less complex at night, where axonal terminals are collapsed together. In contrast, *I_h_* downregulation leads to axonal projections that display little complexity throughout the day, accompanying the reduced PDF levels. Our speculation on why *I_h_* downregulation leaves both, PDF and terminal complexity at levels similar to ZT14 is that I_h_ underlies high activity bursting firing, a property that is functional during the day. Downregulation of this channel impairs this high frequency bursting that would be associated to increased PDF levels and the spreading of sLNv axonal projections in the morning, both phenomena that have been described to be clock and activity-dependent (31, 45, 46). Moreover, we have previously described that structural plasticity depends on PDF levels (47), so the collapsed state of the projections could be linked to PDF decrease as well. To corroborate whether the defects shown upon *I_h_* downregulation are linked to reduced PDF levels we used the GeneSwitch system to express *pdf* in the context of *I_h_* downregulation. **Fig 4A, C-D** shows that indeed, in the context of a surplus of PDF, cycling of this neuropeptide in the sLNvs axonal terminals is restored, while PDF expression in controls cycles with reduced (yet significant) amplitude.

**Fig 4:**
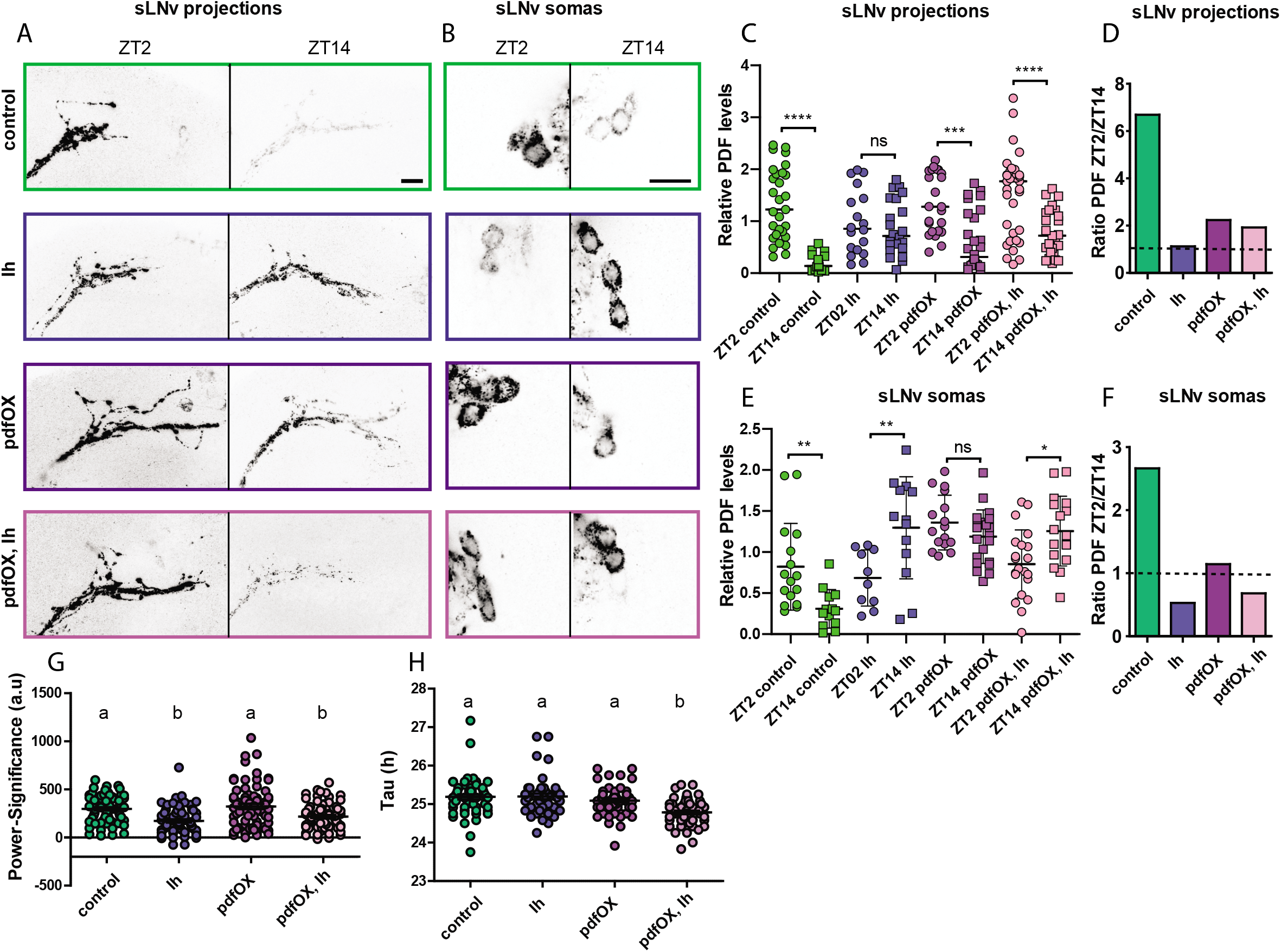
PDF transport is affected upon *I_h_* manipulation. **A, B)** Confocal images of representative sLNv projections (A) and somas (B) of individual flies of *pdf*GS/+ (control), 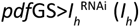, *pdf*GS>UAS-*pdf* (*pdf*OX) and *pdf*GS>UAS-*pdf*, ζ^RNAi^ (*pdf*OX, *I_h_*) at day (left) and night (right) showing their PDF content. Flies were kept in LD 12:12 at 25°C for 7 days in food containing RU486. Brains were dissected at ZT02 and ZT14 and standard anti-PDF immunofluorescence detection was performed, bars indicate 10μm. **C, E)** PDF quantitation of the sLNv dorsal projections (C) or somas (E) for the four genotypes mentioned before. Circles represent day time, squares, night time; each color is a different genotype. Asterisks represent significant statistical differences. For the projections, a non-parametric ANOVA Kruskal-Wallis test and Dunn’s comparisons test showed differences among the two time points in control, *pdfOX* and *pdf*OX, *I_h_* groups but not in *I_h_* group [Kruskal-Wallis statistic (8,196)=71.95, p<0.0001, n=18-28]. Immunoreactivity from somas was analyzed with one-way ANOVA and Sidak’s multiple comparisons test and revealed differences between the two time-points in every genotype except *pdf*OX, although *I_h_* and *pdf*OX, *I_h_* showed differences in the anti-phase direction compared to the control, ANOVA F (7, 120)=10.95, p<0.0001, n=10-22 (each point is the average of 3-4 cell somas for one hemi-brain of an individual fly). **D, F)** Morning to evening PDF level ratios for axonal projections (D) or somas (F). **G)** Locomotor behavior under constant darkness of the same genotypes as before. Experiments were performed as in Fig 1 and Table 2. The rhythmicity measured as power-significance was analyzed by Kruskal-Wallis test followed by Dunn’s comparisons test and showed a significant reduction of power-significance in *I_h_* and *pdf*OX, *I_h_* compared to control and *pdf*OX as indicated by different letters [Kruskal-Wallis statistic (4,31)=31.40, p<0.0001, n=65-72]. **H)** Free running period values were analyzed as well. The same type of analysis reveals a reduction of tau in *pdf*OX, *I_h_* compared to all the other genotypes as indicated by a different letter [Kruskal-Wallis statistic (4,31)=38.28, p<0.0001, n=45-58].

To investigate whether the decreased PDF levels seen at the dorsal projections are due to decreased PDF production or to a failure to recruit PDF-loaded vesicles (i.e. transport) towards the axonal terminal, we measured PDF levels in the sLNv somas. We analyzed somatic PDF levels (see methods) and found that PDF immunoreactivity cycles in the sLNv somas in a way that resembles its cycling at the axonal terminals, with more PDF during the early morning and less PDF at the beginning of the night (**Fig 4B, E-F**). Interestingly, in the context of *I_h_* downregulation, somatic PDF shows an abnormal accumulation during the night, which could be due to a decreased day-time transport towards the axonal terminals that results in anti-phase cycling of somatic PDF levels. PDF overexpression *per se* increases overall levels, preventing PDF cycling in the somas, albeit not in the terminals. On the other hand, PDF overexpression in the context of *I_h_* knock down does not rescue the night-time abnormal PDF accumulation in the somas, however, it does rescue cycling in the projections (**Fig 4C**).

Although PDF overexpression rescues some of the I_h_-related phenotypes at the cellular level, it fails to rescue free running behavior (**Fig 4 G-H**). A plausible explanation for this may be that PDF cycling in the terminals, although rescued, still shows reduced amplitude (**Fig 4D**) and may not be enough to synchronize the remaining clusters. Alternatively, *I_h_* downregulation and the associated reduction of bursting frequency may be affecting the release of other neuropeptides or neurotransmitters besides PDF, which might also contribute to the neuronal communication needed to maintain rhythmicity under constant conditions. PDF expression in the context of *I_h_* downregulation subtly shortens the free-running period (**Fig 4H**), which is reminiscent of reduced PDF levels (2), although the underlying mechanisms remain to be explored.

Overall, these results indicate that I_h_ defines an essential property of the sLNvs that ensures proper regulation of neuropeptide levels and structural plasticity, and provide a causal link between the alteration of electrical activity and the disruption of circadian behavior. Moreover, the careful determination of PDF levels in the sLNv somas suggest that in the context of *I_h_* downregulation there is defective PDF transport towards the axonal projections, underscoring that action potential firing is responsible for an active recruitment of dense core vesicles to the terminals.

### Sleep and the I_h_ channel

We then examined whether reduction in bursting firing frequency and hence, neuropeptide release, could affect sleep behavior. We first quantified sleep behavior in 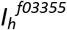 mutants and found that homozygotes displayed an increase in total sleep, mainly due to a significant rise in the number of sleep bouts, which while shorter in duration did not compensate, and resulted in an increase in total sleep during nighttime. Notably, the increase in sleep was more conspicuous towards the end of the night (**Fig 5A-D, Table 3**). Given the ubiquitous nature of this genetic manipulation we reasoned that the deconsolidated sleep phenotype could arise from the lack of *I_h_* in a plethora of neurons; we therefore continued the analysis using *I_h_* RNAi-mediated downregulation in specific LNv clusters.

**Table 3:**
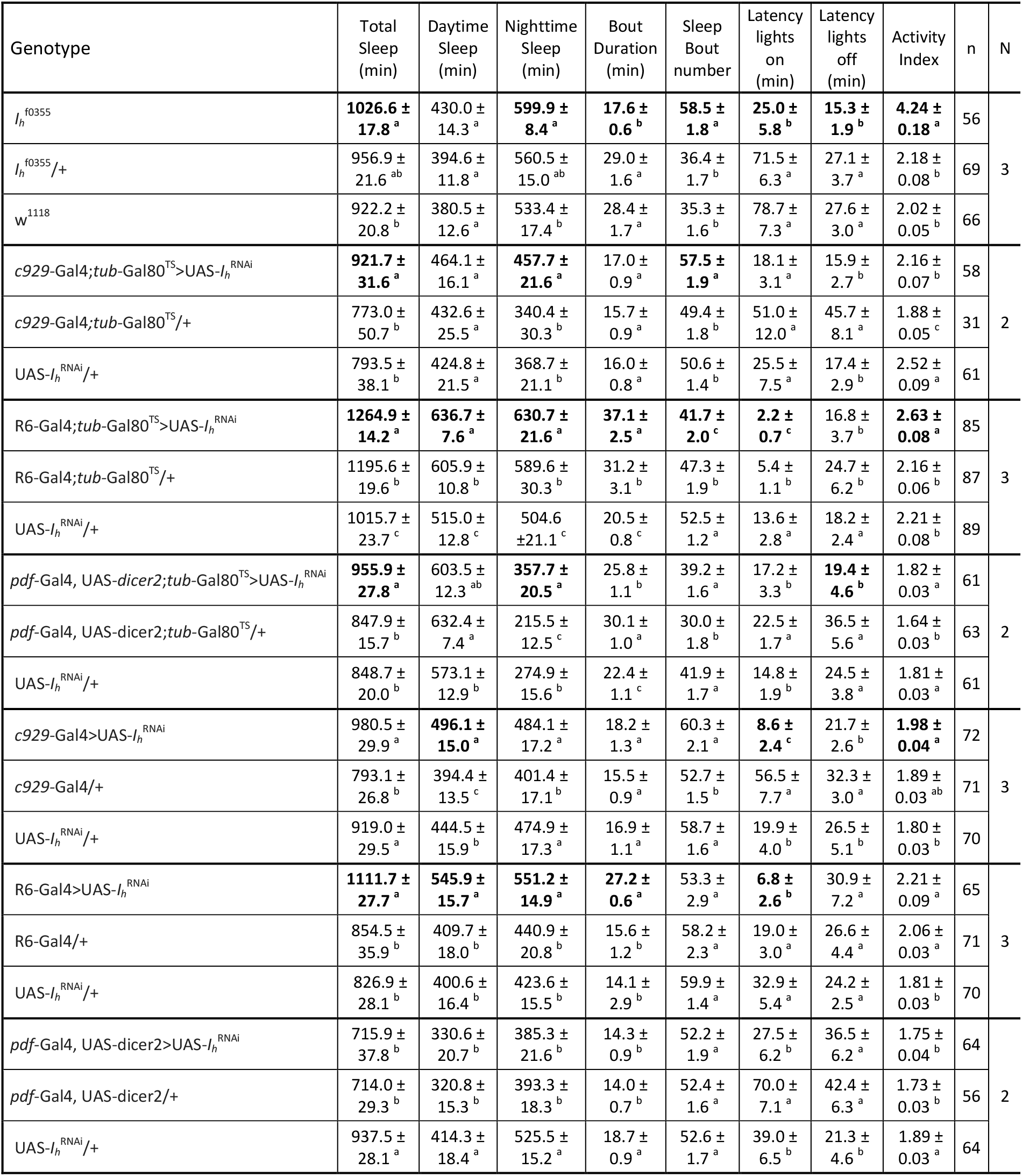
Sleep parameters after genetic manipulation of *I_h_*. The following sleep parameters upon the different genetic manipulations presented in the first column are shown; total sleep, daytime sleep, nighttime sleep, sleep bout duration, bout amount, latency to lights on, latency to lights off, and activity index (defined as the average activity counts in the active minutes). Average ± SEM (Standard Error of the Mean) of N experiments using a final n number of individuals are displayed. Different letters indicate significant differences (p<0.05) after non parametric Kruskal-Wallis statistical analysis with multiple comparisons (p adjustment method= BH). Sleep parameters where the experimental genotype showed statistically significant differences compared to genetic controls are displayed in bold.

**Figure 5:**
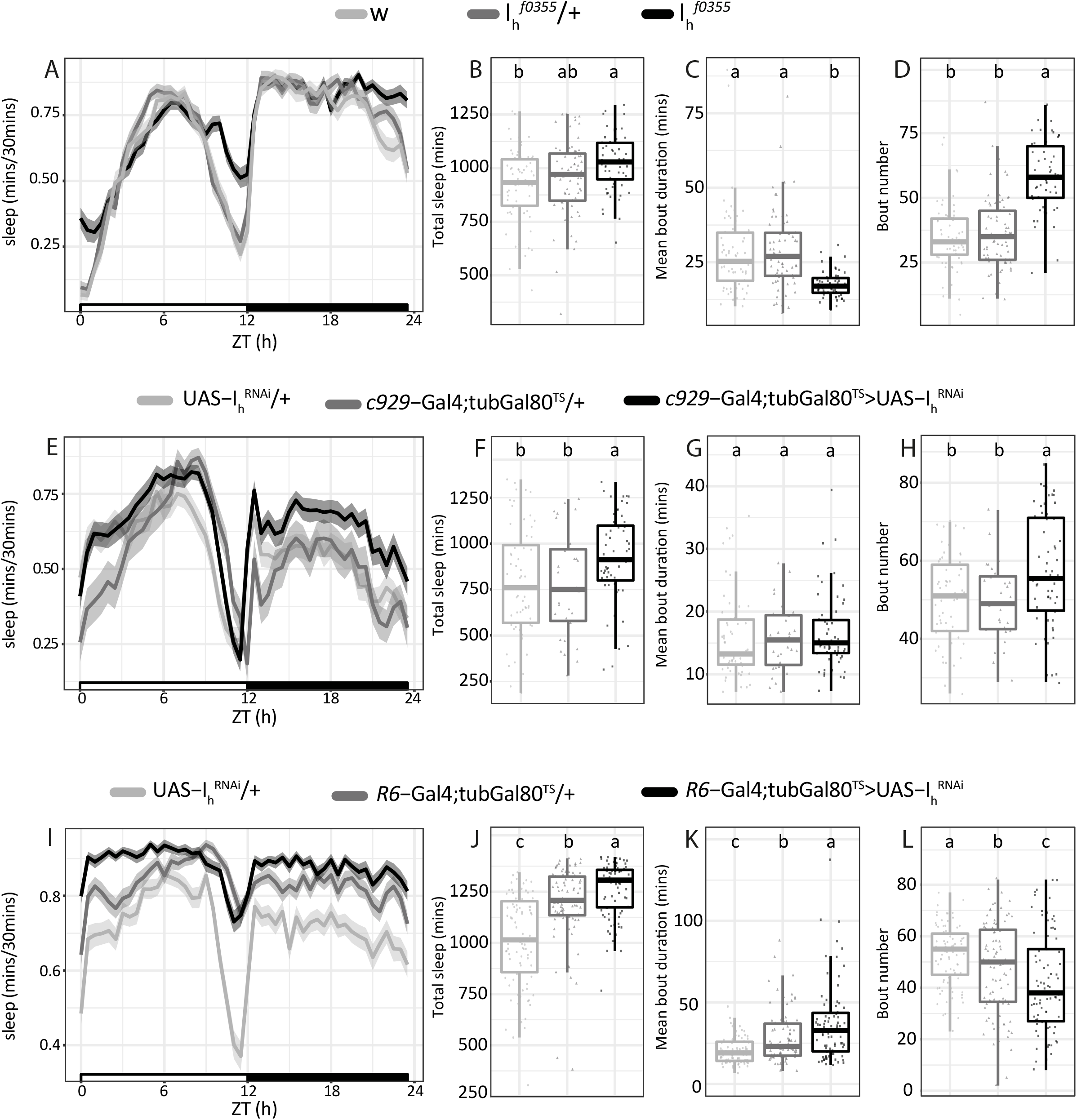
Genetic manipulations of *I_h_* increase sleep. **A, E, I)** Sleep ethograms for the indicated genotypes, quantification of the relative amount of sleep every 30 minutes as a function of the time of the day (starting at zeitgeber time = 0, when lights are turned on) and its standard deviation (shadowed area). Black and white bars at the bottom represent daytime (white) and nighttime (black). **B, F, J)** Boxplots showing the total amount of sleep minutes for each genotype. **C, G, K)** Boxplots showing the average duration of sleep episodes for each genotype. **D, H, L)** Boxplots showing the total amount of sleep episodes for each genotype. For all the boxplots, different letters indicate significant differences (p<0.05) after non parametric Kruskal-Wallis statistical analysis with multiple comparisons (p adjustment method= BH). For more information on sleep parameters see Table 3.

It has previously been demonstrated a significant role of the lLNvs in arousal, as the PDF released by these neurons works as a strong arousal signal (7-9). We therefore analyzed sleep after acute downregulation of the I_h_ channel in the lLNvs and other non-circadian peptidergic neurons (combining the *c929-Gal4* driver with the TARGET system (48)). Similar to the *I_h_* mutants, these flies exhibited an increase in the number of sleep episodes that resulted in a significant rise in nighttime sleep (**Fig 5 E-H, Table 3**); however the duration of the sleep bouts remains unchanged, indicating that the short sleep bout phenotype observed in *I_h_* mutants must derive from the lack of *I_h_* in neurons not covered by the *c929*-Gal4 driver.

Although PDF released from the sLNvs has not been shown to play an arousal role as the one released by the lLNvs, diverse lines of evidence lend support to the notion that sLNvs can have an impact on sleep behavior (49, 50). To test if *I_h_* from sLNvs had any influence on sleep behavior, we resorted to the sLNv-specific driver R6-Gal4 (51). As a consequence of acute downregulation of the I_h_ channel in sLNvs, flies showed a robust increase in the amount of sleep, both at daytime and nighttime (**Fig 5 I-L, Table 3**). Surprisingly, this rise in sleep was due to a more consolidated sleep, as the sleep bout number was reduced but episodes lasted longer in the experimental flies compared to the genetic controls. *I_h_* downregulation experiments were also performed constitutively, showing similar tendencies than the acute ones (**Table 3**).

Given that *I_h_* downregulation in both clusters resulted in an increase in sleep with different characteristics, we tested whether *I_h_* downregulation in all PDF-positive neurons would result in an additive effect. However, this was not the case, and concomitant *I*h knock down correlates with an increase in sleep of similar magnitude. In fact, in case of constitutive downregulation, no significant differences were identified (**Table 3**). Thus, both groups could be contributing to sleep regulation through different mechanisms/signals that, when impaired at the same time, result in a nonlinear combination of effects. Acute and cell-type specific manipulations of LNvs are therefore required to dissect the control of sleep behavior.

Collectively, our work demonstrates that I_h_ certainly plays a role in the control of sleep behavior, both on the overall levels and the timing of sleep. Alterations in the timing of sleep is particularly prevalent, highlighted by a recurrent decrease in the latency to the first sleep episode after lights-on observed in the majority of the *I_h_* genetic manipulations (**Table 3**). Further work will be necessary to pinpoint how different neurons recruit I_h_ to regulate various aspects of their physiology. In particular, the role of neuropeptides in sleep control is widely recognized and involves many neurons throughout the brain. We have initiated here an analysis that includes the sLNvs, the lLNvs+other peptidergic non-circadian neurons and the sLNvs+lLNvs, but it is likely that I_h_ manipulation will impair neuropeptide trafficking in other sleep-related neurons as well.

## Discussion

The physiology of a particular neuron is not regulated by a single ion channel type, but by a complex array of different players; they go from the leak conductances that determine input resistance and resting membrane potential which influence dendritic processes, including summation and propagation of synaptic inputs, to the abundance and quality of voltage-gated ion channels that determine the dynamics of action potential firing, and ultimately dictate the release of classical neurotransmitters and neuropeptides. If we add to this picture the channels that are directly or indirectly activated by ligands we will be able to comprehend, and model, neuronal physiology. We have performed a downregulation screen to describe novel ion channels playing roles in establishing the electrical properties of the LNvs, with the aim of advancing the understanding of LNvs physiology. We focused our attention on I_h_, a poorly studied ionic current in *Drosophila*.

Since the discovery of the first hyperpolarization-gated current in cardiac function (52) a great deal of information has been gained about the role of this type of channels in determining the physiology of the mammalian heart and brain (26). An interesting feature of I_h_ channels is that they are not only sensitive to hyperpolarization, but are also modulated by cyclic nucleotides, hence the name of the channel family HCN (for Hyperpolarization Cyclic Nucleotide-gated). The mammalian genome contains four HCN channel genes HCN1-4, each with specific activation characteristics, distinct but in some cases partially overlapping expression patterns, and different roles in neuronal physiology (53). *Drosophila*’s I_h_ channel is the sole member of the HCN family in its genome (54), but up to 12 different splice variants can be generated, providing diverse channels with particular biophysical properties (55). A phylogenetic analysis indicates that *Drosophila I_h_* (also referred in the literature as DMIH) diverged from a common ancestor before the emergence of the four vertebrate subtypes (56). Interestingly, the domain organization of I_h_ is similar to its vertebrate counterparts, and the interaction between domains is conserved to the point that domain swapping between *Drosophila* I_h_ and vertebrate HCN channels produce similar biophysical results (57).

*Drosophila I_h_* has not been explored in depth yet but it has been reported in the visual system where it regulates the release of glutamate from amacrine cells (33), and at the larval neuromuscular junction where it affects neurotransmitter release (58). An analysis of *I_h_* mutants shows that this channel controls a variety of behaviors (32). Particularly relevant for our work is the fact that *I_h_* has been reported to control circadian rhythms and sleep in *Drosophila* by acting on dopaminergic neurons (59). Although Gonzalo-Gomez *et al*. (2012) did not find any PDF disruptions in the sLNvs of the *I_h_* mutants they generated, our current analysis of adult-specific downregulation of *I_h_* shows that it does affect PDF levels and the structural plasticity at the sLNvs dorsal projections, as well as the accumulation of PDF in the somas albeit with altered circadian dynamics. This highlights the importance of using strategies where a genetic manipulation is performed acutely, in order to avoid homeostatic compensations that may conceal a phenotype. Altogether the collective evidence indicates that *I_h_* may be modulating circadian rhythms and sleep by exerting its role in more than one neuronal type. Whether the molecular mechanisms that regulate, and are regulated by *I_h_* in LNvs and in dopaminergic neurons are similar will require further examination.

Sleep behavior has been previously reported in *I_h_* mutant flies, and, taking into account our contribution, the accumulated evidence raises some controversy that deserves special attention. Using an independently generated *I_h_* null mutant, Gonzalo-Gomez *et al*. (2012) reported that total sleep was unchanged; but they showed, similar to our results, a deconsolidation of sleep resulting from an increase in sleep bout number of shorter duration (59). On the other hand, during the initial characterization of the mutants used in our study, Chen *et al*. reported the opposite sleep phenotype, that is, a decrease in total sleep, with no changes in bout number (32). One important element to take into account is that *I_h_* mutant flies are hyperactive (**Table 3**) and therefore inferring sleep from activity data should be approached with caution. Since *I_h_* mutants display an increase in sleep, and hyperactivity would result in an underestimation of sleep, our results are validated. It is not uncommon to come across published fly sleep data inferred from activity monitoring where basal activity is not reported (32), a practice that warrants further attention.

Perhaps more informative than the mutants is our sleep analysis following acute and cluster-specific downregulation of *I_h_*. We show here that *c929*+ peptidergic neurons use *I_h_* to promote arousal, which in the case of the lLNvs would likely be mediated by PDF release following high frequency neuronal bursting. However, the release of other neuropeptides could also be affected by *I_h_* downregulation; in fact additional neurons besides lLNvs contribute to the waking state within the *c929*-Gal4 driven group (8). Moreover, specific downregulation of *I_h_* in the sLNvs also results in an increase of sleep, however, this increase corresponds to a more consolidated sleep and therefore, although an increase in total sleep is produced upon *I_h_* downregulation with both R6-Gal4 and *c929*-Gal4, the properties of these sleep increase are different. This may indicate that, either the neuropeptide/s these neuronal clusters are releasing with the help of I_h_, or the effects these signals are conveying to their particular postsynaptic targets, are probably different. These findings should be thoroughly characterized in the future, as they provide clear evidence of the participation of the sLNvs in the neuronal circuits governing sleep. Interestingly, in a recent genome-wide association study, ion channels were one of the two main pathways associated to sleep duration both in humans and flies, indicating an evolutionarily conserved function of ion channels in regulating a complex behavior such as sleep (60).

Although no role has been directly demonstrated for I_h_ in *Drosophila* clock neurons before our work, the HCN family has been proposed to contribute to the circadian variations in neuronal excitability in the mammalian suprachiasmatic nucleus (SCN) (61). HCN channels have been reported to be expressed in the SCN (62) but their function has been difficult to discern due to a lack of strong and significant phenotypes following genetic and pharmacologic manipulations, which could be due to the heterogeneity of the SCN neuronal population and the genetic compensation that may arise from having several HCN channel genes (63-65). Taking advantage of the fact that *Drosophila* clock neuron clusters are well identifiable, and that there is only one member of the HCN channel family, we were able to show that I_h_ is a crucial player defining the high activity bursting physiology of LNvs, and that this regulates neuronal outputs and behavior.

The role of cyclic nucleotide cascades in LNvs has been mainly focused on their capacity as transducers of information that impacts on gene expression in the context of the regulation of the circadian clock (66, 67). Our screen has uncovered two cyclic nucleotide-modulated channels (I_h_ and CngA), suggesting that the integration of information signaled by cyclic nucleotides is crucial for circadian function also at rapid time frames, a hypothesis that has already been proposed (67, 68). The case of I_h_, being modulated by both hyperpolarizing voltage and cyclic nucleotides, provides additional complexity, as they could serve as coincidence detectors (scheme in **Fig 6**). The biophysics of I_h_, and therefore the firing properties of LNvs, are affected by both the membrane voltage and the levels of cyclic nucleotides, therefore it is likely that the timing of arrival of these signals may significantly affect the LNvs neuronal output. Albeit purely speculative for the LNvs, the I_h_ current has been proposed to work as a coincidence detector in other systems (69-71). Further research will be necessary to reveal which are the neuronal inputs that contribute to the hyperpolarization and to the variations of cyclic nucleotide levels. Interestingly, HCN channels have been reported to be activated by Vasoactive Intestinal Peptide (72), which is considered a functional homolog of PDF. Therefore, activation of the PDFR signaling cascade could result in an increase in cyclic nucleotide (i.e., cAMP) thus modulating I_h_, adding players to the already complicated integration of synaptic and cell-autonomous cues coordinated at the sLNvs.

**Fig 6:**
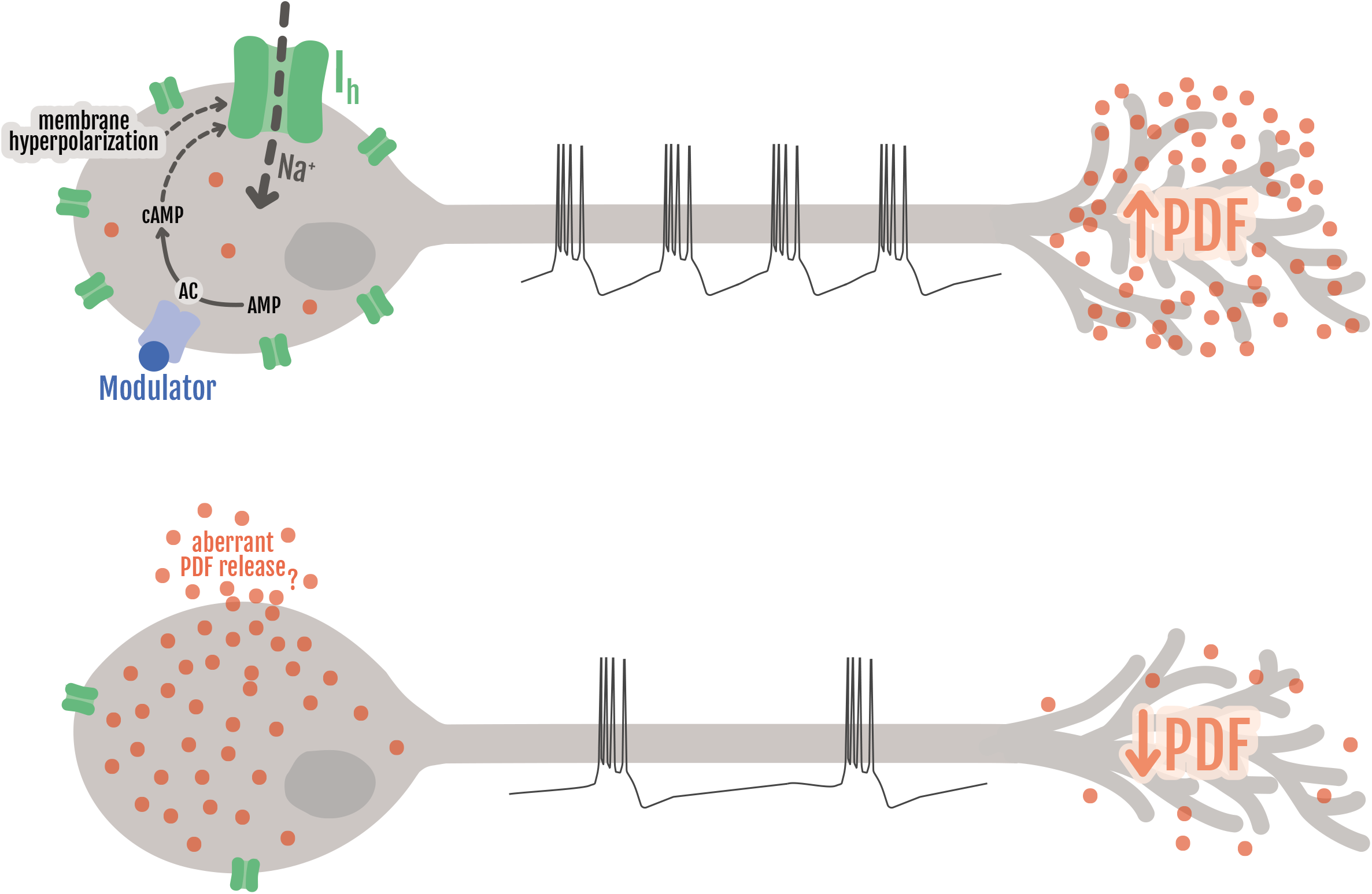
Model summarizing the findings reported and the hypotheses raised by this work. I_h_ channel (in green) responds to membrane hyperpolarization and is modulated by cyclic nucleotides, serving as coincidence detector for electrical and chemical signals mediated by ligands that activate G protein-coupled receptors (such as PDF, Dopamine or other neuropeptides, symbolized by “Modulator”, in blue). I_h_ function is necessary to allow LNvs to fire action potentials in a high frequency bursting mode, which permits the release of PDF (in orange) at high levels and in a timely manner (top neuron). In the absence or upon *I_h_* knock down (bottom neuron), bursting does not reach such high frequency and PDF levels at the axonal projections are reduced. Associated to the decreased bursting frequency, large quantities of PDF accumulate at the soma, likely due to a failure in dense core vesicle transport. This may give rise to a hypothetical aberrant PDF release from the overloaded soma, likely overriding the internal temporal control. All in all, these cellular disruptions result in anomalies at the behavioral level, such as disorganization of circadian locomotor activity and an increase in sleep. At the level of the axonal projections, the model represents an early daytime situation, where in control animals PDF levels are high and the axonal terminal are spread out. However, the accumulation and possible aberrant release of PDF from the soma is more likely to happen during the night (see Fig 4).

One question that remains to be answered is whether the circadian clock is directly regulating I_h_ function in LNvs. Although our work has not focused on this issue, a plausible hypothesis is that I_h_ expression levels may be changing at different times of the day. This is suggested by the work by Abruzzi *et al*. (2011) where they performed chromatin immunoprecipitation tiling array assays with a number of circadian proteins, and showed that the circadian transcription factor CLOCK cycles in its binding to *I_h_* regulatory sequences in *Drosophila* heads (73). Among all the positive hits of our screen, I_h_ is the only one that appears to be directly controlled by the circadian clock according to Abruzzi *et al*. (2011).

Our experiments demonstrate that the sLNvs, considered a central piece of the clock neuron circuitry puzzle, organize their action potential firing in bursts. This bursting frequency depends on synaptic input as it has been shown for the lLNvs (29). LNvs bursting frequency seems to be also influenced by cell autonomous mechanisms since, as we demonstrate here, a null mutation in *I*_h_ produces a decrease in this parameter. Interestingly, the mutation produces a decrease that is of the same magnitude in sLNvs and lLNvs, suggesting that this ion channel regulates bursting frequency in comparable ways in both neuronal types. Remarkably, the DN1 clock neuron cluster has recently been shown to fire action potentials in bursts and that this temporal coding, i.e. the timing of spiking, is relevant for the control of sleep behavior (16).

The importance of gaining as much information as possible about the I_h_ current is underscored by the discovery of several I_h_ channelopathies. Information from both, patients and genetic animal models, has brought to light the relation of mutations on *HCN* channel genes, or accessory subunits, to different conditions such as of epilepsies, autism spectrum disorders, neuropathic pain, Parkinson’s disease, depression and cardiac dysfunction among others (74-76). In this context, learning about *Drosophila* I_h_ helps understanding the basic characteristics of this current and *Drosophila*, with its less complex genome and fantastic genetic amenability, could serve in the future as a model organism to discover interacting proteins and pathways, to ultimately unravel the underlying pathological mechanisms of I_h_ channelopathies.

## Materials and Methods

### Fly Strains

All fly strains used in this study are detailed in **S2 Table**. UAS lines for RNAi downregulation of candidate ion channels were obtained from the Bloomington Stock Center (the ones associated to the *Drosophila* RNAi Screening Center, DRSC), from the Vienna *Drosophila* Resource Center (VDRC) and from the National Institute of Genetics Fly Stock Center (NIG). Information for each of these lines is also available in Table 1 (for the positive hits of the genetic screen) and in S1 Table (for the negative hits). Flies were grown and maintained at 25°C in standard cornmeal medium under 12:12h light:dark cycles unless stated otherwise. For experiments involving the adult-specific GeneSwitch expression system, 2-4 day old adult males raised in normal cornmeal food were transferred to food containing RU486 (mifepristone, Sigma) in 80% ethanol to a final concentration 200 μg/ml or with the same amount of ethanol (vehicle) in control treatments. These experiments were done with a line that includes a UAS-CD8::GFP transgene on the II chromosome. For experiments involving the TARGET system (48) flies were raised at 21°C, induction of the expression system was achieved by increasing the temperature to 30°C. Newly eclosed males were used for all circadian rhythmicity experiments, 3-7 day-old non-virgin females were used for sleep and electrophysiology experiments, a mix of males and females was used for immunofluorescence determination.

### Locomotor Behavior Analysis

Flies were entrained to 12:12h light:dark cycles during their entire development, and newly eclosed adult males were placed in 65 x 5 mm glass tubes and monitored for activity with infrared detectors and a computerized data collection system (Trikinetics, Waltham, MA). For experiments involving the GeneSwitch expression system, newly eclosed adult males were placed in glass tubes containing standard food (supplemented with 200 mg/ml RU486 or vehicle, as indicated) and monitored for activity. Activity was monitored in LD conditions for 3-4 days, followed by constant darkness for at least nine days (DD1–9). Period and rhythmicity parameters as FFT and power were estimated using ClockLab software (Actimetrics). Flies with a single peak over the significance line (p<0.05) in a χ^2^ analysis were scored as rhythmic, which was confirmed by visual inspection of the actograms. For LD anticipatory analysis, the last day before switching to DD was used. Average activity plots at the population level were produced using the Clocklab average activity function for each animal, relativized to its own activity, integrated in 30 minutes bins and then the population average for each genotype was calculated. Morning Anticipation Index (MAI) was calculated as follow, the sum of relativized activity from ZT21.5 from the previous day to ZT0 was divided by the sum of relativized activity from ZT19 from the previous day to ZT0 for each animal. Since data was not normally distributed a non-parametric ANOVA analysis, Kruskal-Wallis test followed by Dunn’s multiple was used to test statistically significant differences. An equivalent procedure was performed for the Evening Anticipation Index (EAI) using data from ZT7 to ZT12.

### Sleep Behavior Analysis

Female flies were socially housed in vials from eclosion at 25°C under 12:12h light:dark cycles until they were 4 to 6 days-old and afterwards transferred to 65 x 5 mm glass tubes (Trikinetics, Waltham, MA) containing normal cornmeal food. Tubes were loaded onto DAM monitors and locomotor activity was assessed using the DAM system under 12:12h light:dark cycles. Sleep data was calculated on the second day after fly loading into tubes to allow them to recover from anesthesia and to acclimate to the new environment. For experiments using the TARGET system (77) flies were raised at 21°C, socially housed in vials from eclosion until they were 6 days-old and afterwards transferred to 65 x 5 mm glass tubes. Monitors were kept for two days at 21°C to measure sleep under the restrictive temperature at which the RNAi is not expressed (which in all cases produced no effect), and then the incubator temperature was raised to 30°C for two more days to allow RNAi expression, always under 12:12h light:dark cycles. Sleep data was calculated on the second day at 30°C. The DAM System binning time was set to 1min. Sleep was defined as no movement for 5min (78, 79). Rethomics, a collection of packages running in R language (80), was used to infer sleep from locomotor activity data, to build graphs of sleep for 30min as a function of the time of day, to get measurements of total sleep, day sleep, night sleep, sleep bout duration, sleep bout number, latencies to lights on and off and to get an activity index (defined as the average activity counts in the active minutes) of each individual fly. Behavioral experiments were conducted at least 2-3 times, with 15 to 30 individuals per genotype.

### Electrophysiology

Three to seven days-old female flies were anesthetized with a brief incubation of the vial on ice, brain dissection was performed on external recording solution which consisted of (in mM): 101 NaCl, 3 KCl, 1 CaCl_2_, 4 MgCl_2_, 1.25 NaH_2_PO_4_, 5 glucose, and 20.7 NaHCO_3_, pH 7.2, with an osmolarity of 250 mmol/kg (based on solution used by Cao and Nitabach, 2008). After removal of the proboscis, air sacks and head cuticle, the brain was routinely glued ventral side up to a sylgard-coated coverslip using a few μls of tissue adhesive 3M Vetbond. The time from anesthesia to the establishment of the first successful recording was approximately 15-19min spent as following: 5–6min for the dissection, 4-5min for the protease treatment to remove the brain’s superficial glia and 6-8min to fill and load the recording electrode onto the pipette holder, approach the cell, achieve the gigaohm seal and open the cell into whole-cell configuration to start recording. LNvs were visualized by red fluorescence in *pdf-RFP* using a Leica DM LFS upright microscope with 63X water-immersion lens and TK-LED illumination system (TOLKET S.R.L, Argentina). Once the fluorescent cells were identified, cells were visualized under IR-DIC using a Hamamatsu ORCA-ER camera and Micro Manager software. lLNvs were distinguished from sLNvs by their size and anatomical position. To allow the access of the recording electrode, the superficial glia directly adjacent to LNvs somas was locally digested with protease XIV solution (10mg/ml, SIGMA-ALDRICH P5147) dissolved in external recording solution. This was achieved using a large opened tip (approximately 20μm) glass capillary (pulled from glass of the type GC100TF-10; Harvard Apparatus, UK) and gentle massage of the superficial glia with mouth suction to render the underling cell bodies accessible for the recording electrode with minimum disruption of the neuronal circuits. After this procedure, protease solution was quickly washed by perfusion of external solution. Recordings were performed using thick-walled borosilicate glass pipettes (GC100F-10; Harvard Apparatus, UK) pulled to 6-7 MΩ using a horizontal puller P-1000 (Sutter Instruments, US) and fire polished to 9-12 MΩ. Recordings were made using an Axopatch 200B amplifier controlled by pClamp 9.0 software via a Digidata 1322A analog-to-digital converter (Molecular Devices, US). Recording pipettes were filled with internal solution containing (in mM): 102 potassium gluconate, 17 NaCl, 0.085 CaCl_2_, 0.94 EGTA and 8.5 HEPES, pH 7.2 with an osmolarity of 235 mmol/kg (based on the solution employed by Cao and Nitabach, 2008). Cell-attached configuration was achieved by gentle suction and recordings were performed in voltage-clamp mode with no hold. For whole-cell configuration, gigaohm seals were accomplished using minimal suction followed by break-in into whole-cell configuration using gentle suction in voltage-clamp mode with a holding voltage of −60mV. Gain of the amplifier was set to 1 during recordings and a 5kHz Lowpass Bessel filter was applied throughout. Spontaneous firing was recorded in current clamp (I=0) mode. Analysis of traces was carried out using Clampfit 10.4 software. Bursting frequency was calculated as the number of bursts in a minute of recording. For comparisons, all recordings were quantified at the same time post-dissection as specified in the text and figure legends. For AP firing rate calculation the event detection tool of Clampfit 10.4 was used. In many cases we were able to see the two different AP sizes reported previously (28), however, for AP firing rate calculation only the large APs were taken into account. Traces shown in figures were filtered offline using a lowpass boxcar filter with smoothing points set to 9. Perfusion of external saline in the recording chamber was achieved using a peristaltic pump (MasterFlex C/L). All recordings were performed during the light phase, between ZT1 and ZT10.

### Immunofluorescence detection

Heads were cut at *zeitgeber* times 2 and 14, fixed in paraformaldehyde 4% in PB 0.1M for 35-45 minutes at room temperature and brains were dissected afterwards, washed 5 times in PBS-Triton X-100 0.1%, blocked with 7% normal goat serum for 2 hours at RT and incubated with primary antibody (see antibodies information in **S2 Table**), ON at 4ºC. After five 15 minutes washes in PBS-Triton X-100 0.1%, brains were incubated with the secondary antibody. Confocal images were obtained in a Zeiss 710 Confocal Microscope or Pascal Confocal Microscope. All the photographs within the same experiment were taken with the same confocal parameters. In the *pdf* overexpression experiments (Fig 4), data was relativized to the average of intensities for each experiment because two different microscopes were used. The acquisition of sLNv soma images required different confocal parameters (laser intensity, gain, zoom).

### PDF quantitation

For the quantitation of PDF intensity at the sLNv projections, we assembled a maximum intensity z-stack that contains the whole projection (approximate 10 images) and constructed a threshold image to create a ROI for measure immunoreactivity intensity using ImageJ (NIH). Data was analyzed with InfoStat software (Universidad Nacional de Córdoba, Argentina) and GraphPad. For quantitation of PDF intensity at the sLNv somas we used a unique 1 μm image per cell, which was the one where the PDF cytoplasm immunoreactivity signal could be clearly differentiated from the empty nucleus. The draw tool from ImageJ (NIH) enabled to measure only the PDF signal at the cytoplasm, and this procedure was repeated for each cell (3-4) in each brain (only one brain hemisphere). Background intensity was subtracted for each brain and average intensity was calculated. Data was normalized using the average intensity for the whole population of brains of the experiment. This way of quantifying PDF in the sLNvs somas allowed a more precise assessment of neuropeptide levels and it may be the reason why we were able to detect circadian cycling of PDF levels, unlike previous reports that were unable to detect them (42). Statistics analysis was done using the GraphPad program, after testing data normality one-way ANOVA and Sidak’s multiple comparisons tests were performed to determine time of -day- genotype differences.

### Analysis of structural plasticity

To assess the degree of complexity within the sLNvs dorsal projections we performed immunofluorescence against a membrane version of GFP. The maximum intensity z-stack image was transformed into a threshold image and Sholl analysis was performed with ImageJ (NIH) software. Each picture was corroborated by visual inspection to confirm the number of crosses in every 10 μm-concentric Sholl ring. Data was analyzed by means of InfoStat software (Universidad Nacional de Córdoba, Argentina).

### Statistical Analysis

The following statistical analyses were used in this study: one-way ANOVA and two-way ANOVA with post hoc Tukey’s HSD test for multiple comparisons of parametric data, and non-parametric Kruskal-Wallis statistical analysis with multiple comparisons (p adjustment method = BH) as specified in figure legends. Parametric tests were used when data were normally distributed and showed homogeneity of variance, tested by Kolmogorov Smirnov test and Levene’s test, respectively. Sidak’s and Dunn’s multiple comparisons tests were performed after parametric and non-parametric ANOVA when GraphPad software was used. Sleep data tended to not show a normal distribution, hence non-parametric statistics were used. Statistical analyses were performed using Infostat for circadian rhythmicity and immunofluorescence experiments, R-based Rethomics package for sleep data and Origin software for electrophysiological parameters. A p value < 0.05 was considered statistically significant. Throughout the manuscript n represents the total number of measurements compared in each experimental group (behavior of an individual, brain morphology, or neuronal recordings, depending of the experiment), and N represents the number of independent times an experiment was repeated. Boxes in box and whisker plots for sleep and electrophysiological parameters represent the median and interquartile range (the distance between the first and third quartiles). In all tables, parameters represent the mean value ± standard error of the mean. In dot plots for circadian power and tau and in fluorescence and structural plasticity quantification lines represent the mean value; error bars depict the standard error of the mean.

## Acknowledgements

M.F.C., N.I.M. and L.F. are members of the Argentine Research Council for Science and Technology (CONICET). This work was supported by the Agencia Nacional de Promoción Científica y Tecnológica (Grants PICT-2011-2185 to M.F.C. and PICT-2011-2364 and PICT-2015-2557 to N.I.M.), the CONICET (Grant PIP-11220130100378 to N.I.M.) and FOCEM-Mercosur (COF 03/11 to IBioBA). The funders had no role in study design, data collection and analysis, decision to publish, or preparation of the manuscript.Stocks obtained from the Bloomington Drosophila Stock Center (NIH P40OD018537), the NIG-Fly Stock Center and the Vienna Drosophila Resource Center (81) were used in this study. We also thank Dr. Zuo Ren Wang for *I_h_* mutant fly stocks. We thank Esteban Beckwith and Quentin Geissmann for indispensable help with sleep quantification software.

