## Supplemental Table 1 for "High frequency neuronal bursting is essential for circadian and sleep behaviors in *Drosophila*"

| Gene | CG | RNAi info | Channel type | n | N |
|---|---|---|---|---|---|
| *Atpα* | CG5670 | DRSC 28073 | Na+/K+ ATPase α subunit | 16 | 1 |
| *Atpα* | CG5670 | VDRC KK 100619 | Na+/K+ ATPase α subunit | 45 | 3 |
| *Calx* | CG5685 | DRSC 28306 | Ca++ Na+ antiporter | 8 | 1 |
| *Calx* | CG5685 | VDRC KK 104789 | Ca++ Na+ antiporter | 13 | 1 |
| *Ca-α1D* | CG4894 | DRSC 25830 | VG Ca++ channel | 16 | 1 |
| *Ca-α1D* | CG4894 | VDRC GD 51490 | VG Ca++ channel | 16 | 1 |
| *Ca-α1T* | CG15899 | VDRC GD 31963 | VG Ca++ channel | 70 | 3 |
| *Ca-α1T* | CG15899 | DRSC 26251 | VG Ca++ channel | 50 | 2 |
| *eag* | CG10952 | DRSC 31678 | VG cation channel | 16 | 1 |
| *eag* | CG10952 | VDRC KK 100260 | VG cation channel | 16 | 1 |
| *Hk* | CG43388 | DRSC 28330 | VG K+ channel β subunit | 16 | 1 |
| *Hk* | CG43388 | VDRC KK 101402 | VG K+ channel β subunit | 14 | 1 |
| *inx2* | CG4590 | DRSC 29306 | Gap junction channel | 13 | 1 |
| *inx2* | CG4590 | VDRC KK 102194 | Gap junction channel | 9 | 1 |
| *Ir* | CG6747 | DRSC 25823 | VG K+ channel | 32 | 2 |
| *Ir* | CG6747 | VDRC KK 107389 | VG K+ channel | 15 | 1 |
| *Irk2* | CG4370 | DRSC 25820 | Inwardly rectifying K+ channel | 31 | 2 |
| *Irk2* | CG4370 | VDRC GD 4341 | Inwardly rectifying K+ channel | 13 | 1 |
| *KCNQ* | CG33135 | DRSC 27252 | VG K+ channel | 65 | 4 |
| *KCNQ* | CG33135 | VDRC KK 106655 | VG K+ channel | 15 | 1 |
| *Ncc69* | CG4357 | DRSC 28682 | Na+ K+ Cl- symporter | 16 | 1 |
| *Ncc69* | CG4357 | VDRC KK 106499 | Na+ K+ Cl- symporter | 16 | 1 |
| *Nckx30C* | CG18660 | DRSC 27246 | Na+ K+ Ca++ exchanger | 15 | 1 |
| *nrv1* | CG9258 | VDRC GD 46542 | Na+/K+ ATPase β subunit | 14 | 1 |
| *nrv1* | CG9258 | VDRC KK 103702 | Na+/K+ ATPase β subunit | 15 | 1 |
| *nrv2* | CG9261 | DRSC 28666 | Na+/K+ ATPase β subunit | 15 | 1 |
| *nrv2* | CG9261 | VDRC GD 2660 | Na+/K+ ATPase β subunit | 23 | 2 |
| *para* | CG9907 | VDRC GD 6131 | VG Na+ channel | 16 | 1 |
| *para* | CG9907 | VDRC KK 104775 | VG Na+ channel | 28 | 2 |
| *picot* | CG8098 | DRSC 25920 | Phosphate Na+ symporter | 15 | 1 |
| *picot* | CG8098 | VDRC KK 101082 | Phosphate Na+ symporter | 14 | 1 |
| *ppk* | CG3478 | DRSC 29571 | Amiloride sensitive Na+ channel | 16 | 1 |
| *ppk* | CG3478 | VDRC KK 108683 | Amiloride sensitive Na+ channel | 34 | 3 |
| *ppk12* | CG10972 | DRSC 27092 | Amiloride sensitive Na+ channel | 16 | 1 |
| *ppk12* | CG10972 | VDRC KK 105131 | Amiloride sensitive Na+ channel | 15 | 1 |
| *ppk25* | CG33349 | DRSC 27088 | Amiloride sensitive Na+ channel | 16 | 1 |
| *ppk25* | CG33349 | VDRC KK 101808 | Amiloride sensitive Na+ channel | 16 | 1 |
| *ppk28* | CG4805 | DRSC 31878 | Amiloride sensitive Na+ channel | 16 | 1 |
| *ppk28* | CG4805 | VDRC KK 100946 | Amiloride sensitive Na+ channel | 12 | 1 |
| *sei* | CG3182 | DRSC 31681 | VG K+ channel | 15 | 1 |
| *sei* | CG3182 | VDRC KK 104698 | VG K+ channel | 16 | 1 |
| *Sh* | CG12348 | DRSC 31680 | VG K+ channel | 16 | 1 |
| *Sh* | CG12348 | VDRC KK 104474 | VG K+ channel | 31 | 2 |
| *Shal* | CG9262 | DRSC 31879 | VG K+ channel | 15 | 1 |
| *Shaw* | CG2822 | DRSC 28346 | VG K+ channel | 16 | 1 |
| *Shaw* | CG2822 | VDRC KK 110589 | VG K+ channel | 16 | 1 |
| *SK* | CG10706 | DRSC 27238 | Ca++ activated K+ channel | 16 | 1 |
| *SK* | CG10706 | VDRC KK 103985 | Ca++ activated K+ channel | 16 | 1 |
| SLO2 | CG42732 | DRSC 32034 | Na+ activated K+ channel | 16 | 1 |
| *stj* | CG12295 | DRSC 25807 | VG Ca++ channel | 15 | 1 |
| *stj* | CG12295 | VDRC KK 108569 | VG Ca++ channel | 15 | 1 |
| *Teh2* | CG15004 | VDRC GD 9037 | VG Na+ auxiliary subunit | 16 | 1 |
| *Teh2* | CG15004 | VDRC KK 104951 | VG Na+ auxiliary subunit | 16 | 1 |
| *Teh4* | CG15003 | VDRC GD 11621 | VG Na+ auxiliary subunit | 13 | 1 |
| *Teh4* | CG15003 | VDRC KK 102161 | VG Na+ auxiliary subunit | 16 | 1 |
| *trp* | CG7875 | DRSC 31649 | Light activated Ca++ channel | 10 | 1 |
| *trp* | CG7875 | VDRC GD 1366 | Light activated Ca++ channel | 14 | 1 |

**S1 Table: Negative hits of the ion channel downregulation behavioral screen.**

This table includes the list of UAS-RNAi transgenic lines that, when driven exclusively in LNvs, did not produced statistically significant alterations in free running period and/or percentage of rhythmicity compared to pdf,dicer/+ control genotype (after Student's T-test analysis). N indicates number of independent experiments performed. n indicates number of individuals tested. The appearance of a gene in this table suggests that it may not be involved in the circadian function of LNvs. However, most of these RNAi constructs have not been individually tested for their actual performance on ion channel knock down. Moreover, it should be noticed that for some genes, such as Shal and Ca-α1T, one RNAi construct was able to affect behavior, while others were not. Further investigations are necessary to determine the roles of these channels in LNvs function. Besides the efficiency of the particular RNAi transgenic line, another phenomenon that should be taken into account is that, in some cases, a homeostatic compensation of ion channel downregulation might have taken place. For instance, it is surprising that targeting the gene coding for the only classical voltage-gated sodium channel paralytic (*para*) in LNvs has not resulted in a behavioral phenotype. Most likely, this genetic manipulation has produced compensation, as it has been reported to happen for such an important and therefore highly regulated ion conductance (82). Interestingly, downregulation of *para* accessory subunit *tipE* does affect circadian behavior (see Table 1), indicating that less compensatory mechanisms may be in place to counterbalance such genetic manipulation, and that affecting *para* in this indirect way is probably having a detrimental effect on LNvs ability to fire action potentials. For all these reasons, this table only provides the information that the specific RNAi transgenic lines shown, in the particular conditions we have used, are not able to affect circadian behavior when driven in LNvs. Further analysis is necessary to make stronger statements in all cases. V: Voltage, G: Gated, DRSC: *Drosophila* RNAi Screening Center, VDRC: Vienna *Drosophila* Resource Center.
