## Supplemental Figure 1 for "High frequency neuronal bursting is essential for circadian and sleep behaviors in *Drosophila*"

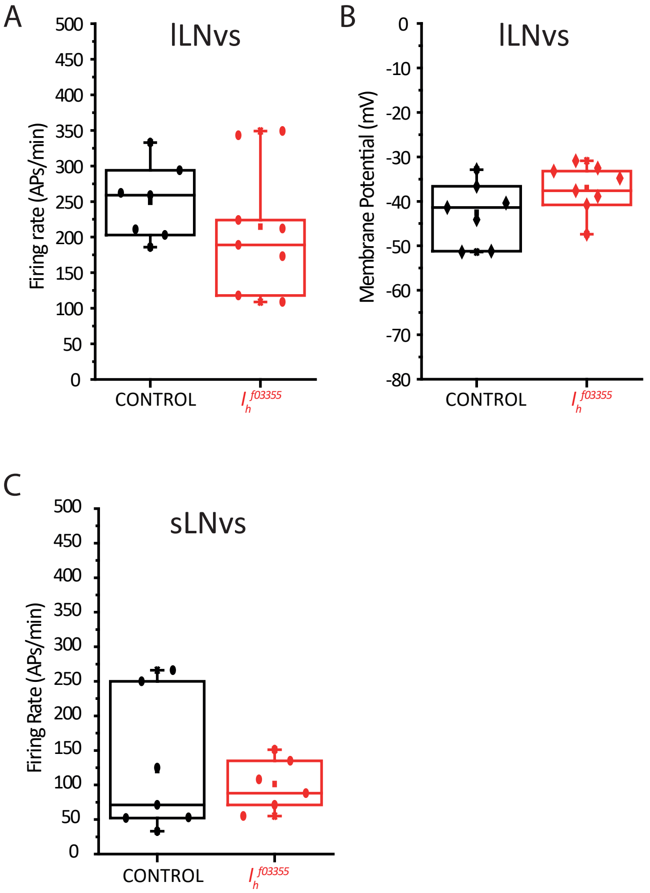


**S1 Fig: Mutation of $I_h$ does not significantly affect other electrophysiological parameters of LNvs.**

**A)** No statistical significant differences were found in action potential firing rate of lLNvs when comparing control (*pdf*-RFP) and $I_h$ homozygote mutant genotypes ($I_h^{f03355}$; *pdf*-RFP). **B)** No statistical significant differences were found in membrane potential (measured as the trough between bursts) of lLNvs when comparing control (*pdf*-RFP) and $I_h$ homozygote mutant genotypes ($I_h^{f03355}$; *pdf*-RFP). **C)** No statistical significant differences were found in action potential firing rate of sLNvs when comparing control (*pdf*-RFP) and $I_h$ homozygote mutant genotypes ($I_h^{f03355}$; *pdf*-RFP). Membrane potential was not quantified in sLNvs as recordings were made in cell-attached configuration and it is not possible to measure this parameter under this configuration. All quantifications were done at exactly 23min post-dissection. In all cases p>0.05 after Student's t-test. n: $lLNvs_{CONTROL}$=7, $lLNvs_{Ihf03355}$=8, $sLNvs_{CONTROL}$=7, $sLNvs_{Ihf03355}$=6.
