## Supplemental Figure 2 for "High frequency neuronal bursting is essential for circadian and sleep behaviors in *Drosophila*"

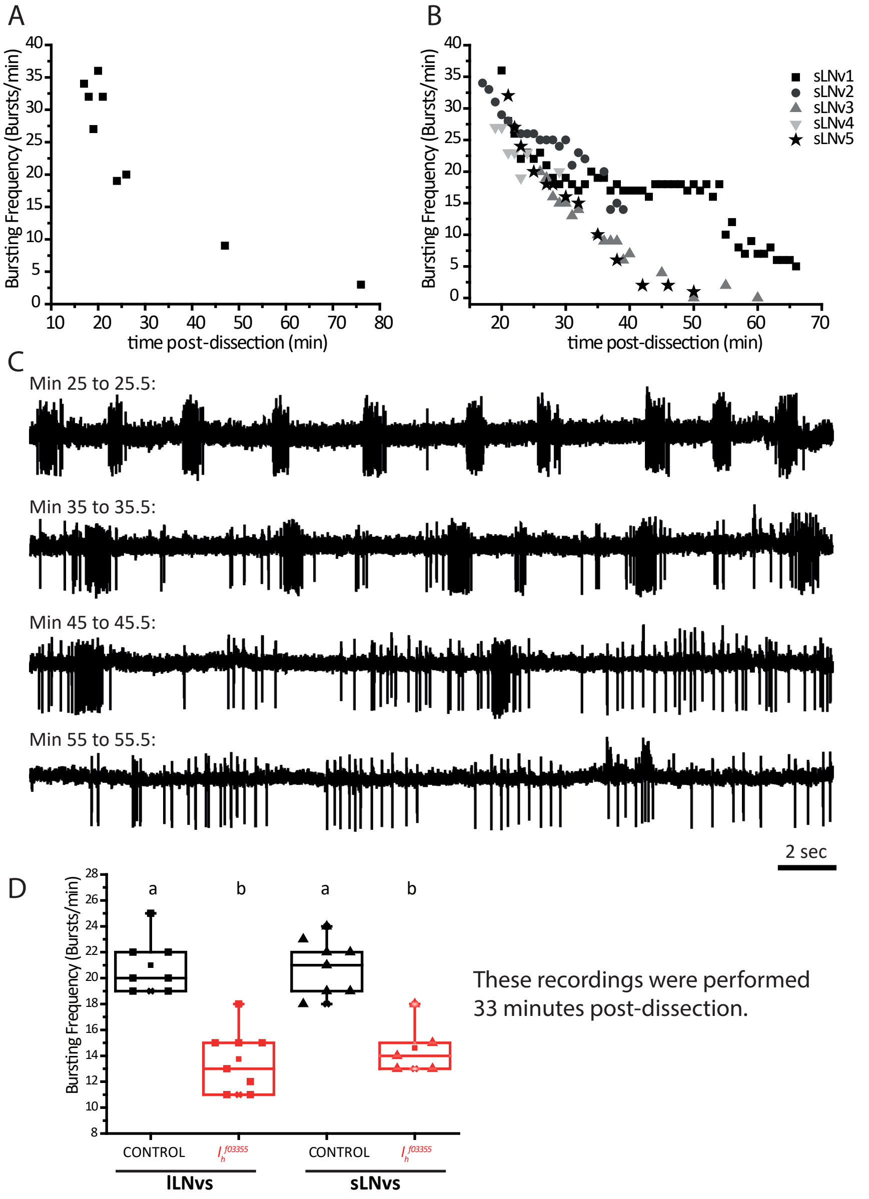


**S2 Fig: sLNvs bursting depends on synaptic inputs.**

As has been demonstrated before for lLNvs (29) we show here that sLNvs bursting frequency also decays as a function of the time ex vivo. **A)** The number of bursts in the initial minute of recording of 9 individual control (*pdf*-RFP) sLNvs recorded at different times post-dissection is shown. For the late points the preparation was left in the chamber on purpose before establishing the recording. **B)** Shows the bursting frequency of 5 individual control (*pdf*-RFP) sLNvs where the recordings were long enough to appreciate the decay in this parameter as a function of time post-dissection not only as a population as in A, but as individual cells. **C)** Shows 30 second windows of cell-attached recording of a representative sLNv (sLNv3 in B) at different times post-dissection. From top to bottom the 30 seconds starting at minutes 25, 35, 45 and 55 post-dissection are shown. At the beginning of the recording, all action potentials are organized in bursts. As time passes, action potentials become less organized in bursts, going through a phase of bursting-tonic firing and becoming purely tonic towards the end. This figure shows that the fact that in A and B the neurons have a tendency towards the cero bursting frequency does not mean that the neurons are not firing, but that they are doing so in a tonic mode. **D)** Box plot showing bursting frequency quantification of lLNvs and sLNvs of control (*pdf*-RFP) and Ih homozygote mutant genotypes ($I_h^{f03355}$; *pdf*-RFP), this quantifications were done at exactly 33min post-dissection. Different letters indicate significant differences ($p<0.05$) after a one-way ANOVA with Tukey test for means comparisons. n: $\text{lLNvs}_{\text{CONTROL}}$=7, $\text{lLNvs}_{\text{Ihf03355}}$=8, $\text{sLNvs}_{\text{CONTROL}}$=8, $\text{sLNvs}_{\text{Ihf03355}}$=5.
