## Supplemental Table 2 for "High frequency neuronal bursting is essential for circadian and sleep behaviors in *Drosophila*"

| REAGENT or RESOURCE | SOURCE | IDENTIFIER |
|---|---|---|
| **Antibodies** | | |
| Rat polyclonal anti-PDF, dilution 1:250 | Depetris-Chauvin et al., 2011 | Not available |
| Chicken polyclonal anti-GFP, dilution 1:250 | Aves Lab | Cat# GFP-1020, RRID:AB_10000240 |
| Alexa Fluor 647-AffiniPure Donkey Anti-Rat, dilution 1:500 | Jackson ImmunoResearch Lab | Cat# 712-605-150, RRID:AB_2340693 |
| Cy2-AffiniPure Donkey Anti-Chicken, dilution 1:500 | Jackson ImmunoResearch Lab | Cat# 703-225-155, RRID:AB_2340370 |
| **Chemicals** | | |
| NaCl, Sodium chloride | Sigma-Aldrich | S7653; CAS: 7647-14-5 (BioXtra, ≥99.5% (AT)) |
| KCl, Potassium chloride | Sigma-Aldrich | P3911; CAS: 7447-40-7 (ACS reagent, 99.0 100.5%) |
| $CaCl_2.2H_2O$, Calcium chloride dihydrate | Sigma-Aldrich | 223506; CAS: 10035-04-8 (ACS reagent, ≥99%) |
| $MgCl_2.6H_2O$, Magnesium chloride hexahydrate | Sigma-Aldrich | M2670; CAS: 7791-18-6 (BioXtra, ≥99.0%) |
| $NaH_2PO_4$, Sodium phosphate monobasic | Sigma-Aldrich | S8282; CAS: 7558-80-7 (BioXtra, ≥99.0%) |
| $NaHCO_3$, Sodium bicarbonate | Sigma-Aldrich | S6297; CAS: 144-55-8 (BioXtra, 99.5-100.5%) |
| D-(+)-Glucose | Sigma-Aldrich | G8270; CAS: 50-99-7 (≥99.5% (GC)) |
| Protease from Streptomyces griseus | Sigma-Aldrich | P5147; CAS: 9036-06-0 (Type XIV, ≥3.5 units/mg solid, powder) |
| Potassium D-gluconate | Sigma-Aldrich | G4500; CAS: 299-27-4 (≥99%) |
| EGTA | Sigma-Aldrich | E3889; CAS: 67-42-5 (for molecular biology, ≥97.0%) |
| HEPES | Sigma-Aldrich | H3375; CAS: 7365-45-9 (≥99.5% (titration)) |
| RU-486, Mifepristone | Sigma-Aldrich | M8046, CAS: 84371-65-3 (≥98%) |
| Paraformaldehyde | Sigma-Aldrich | 441244; CAS: 30525-89-4 |
| NaCl, Sodium chloride (for PBS solution) | Cicarelli Laboratorios | 750; CAS: 7647-14-5 |
| $Na_2HPO_4$, Sodium phosphate dibasic | Sigma-Aldrich | S3264; CAS: 7558-79-4 (for molecular biology, ≥98.5% (titration)) |
| Triton™ X-100 | Sigma-Aldrich | T9284; CAS: 9002-93-1 (BioXtra) |
| Goat serum | Natocor | 734 |
| Vetbond Tissue Adhesive | 3M | 1469SB |
| **Organisms/Strains** | | |
| *D. melanogaster* : *pdf* -GAL: y[1] w[*]; P{w[+mC]=Pdf GAL4.P2.4}2 | Bloomington Drosophila Stock Center | BDSC: previously 6900, now available as part of 25031; FlyBase: FBtp0011844 |
| *D. melanogaster* : UAS-CD8::GFP: y[1] w[*]; P{w[+mC]=UAS-mCD8::GFP.L}LL5, P{UAS-mCD8::GFP.L}2 | Bloomington Drosophila Stock Center | BDSC: 5137; FlyBase: FBst0005137 |
| *D. melanogaster* : *c929* -Gal4: w[*]; P{w[+mW.hs]=GawB}dimm[929] crc[929] | Bloomington Drosophila Stock Center | BDSC:25373; FlyBase: FBst0025373 |
| *D. melanogaster* : tub-Gal80$^{TS}$: w[*]; P{w[+mC]=tubP-GAL80[ts]}2/TM2 | Bloomington Drosophila Stock Center | BDSC: 7017; FlyBase: FBst0007017 |
| *D. melanogaster* : *pdf* -GeneSwitch: w*; P{UAS-mCD8::GFP.L}LL5; P{Pdf-GS}3/TM3, Sb1 | Bloomington Drosophila Stock Center, Depetris-Chauvin et al., 2011 | BDSC: 80956; FlyBase: FBst0080956 |
| *D. melanogaster* : UAS-*dicer2* : w[1118]; P{UAS-dicer2, w[+]} | Vienna *Drosophila* Resource Center | VDRC ID: 60008 |
| *D. melanogaster* : R6-Gal4: P{?GawB}crc$^{R6}$ | Helfrich-Forster et al., 2007 | FlyBase: FBti0016844 |
| *D. melanogaster* : *pdf-RFP* : Pdf-RFP transgene has 0.6 kb of Pdf regulatory genomic DNA 0.5 kb upstream the start site of transcription and 0.1 kb downstream) fused to DNA encoding mRFP1, a monomeric soluble red fluorescent protein (Shaner et al., 2004). Injected into *y* w flies. | Reuben et al., 2012 | FlyBase: FBrf0219602 |

| | | |
|---|---|---|
| *D. melanogaster* : $I_h^{f01485}$ : PBac{WH}Ihf01485 | Exelixis at Harvard Medical School | FlyBase: FBst1017022 |
| *D. melanogaster* : $I_h^{f03355}$ : PBac{WH}Ihf03355 | Exelixis at Harvard Medical School | FlyBase: FBst1018427 |
| *D. melanogaster* : RNAi of *cac* : P{KK101478}VIE-260B | Vienna *Drosophila* Resource Center | VDRC ID: 104168; FlyBase: FBst0476026 |
| *D. melanogaster* : RNAi of *cac* : y[1] v[1]; P{y[+t7.7] v[+t1.8]=TRiP.JF02572}attP2 | Bloomington Drosophila Stock Center | BDSC: 27244; FlyBase: FBst0027244 |
| *D. melanogaster* : RNAi of *Ca-α1T* : P{KK100082}VIE-260B | Vienna *Drosophila* Resource Center | VDRC ID: 108827; FlyBase: FBst0480621 |
| *D. melanogaster* : RNAi of *ClC-a* : P{KK101247}VIE-260B | Vienna *Drosophila* Resource Center | VDRC ID: 110394; FlyBase: FBst0481966 |
| *D. melanogaster* : RNAi of *CngA* : y[1] v[1]; P{y[+t7.7] v[+t1.8]=TRiP.JF02039}attP2 | Bloomington Drosophila Stock Center | BDSC: 26014; FlyBase: FBst0026014 |
| *D. melanogaster* : RNAi of *CngA* : P{KK108314}VIE-260B | Vienna *Drosophila* Resource Center | VDRC ID: 101745; FlyBase: FBst0473618 |
| *D. melanogaster* : RNAi of $I_h$ : P{KK100190}VIE-260B | Vienna *Drosophila* Resource Center | VDRC ID: 110274; FlyBase: FBst0481852 |
| *D. melanogaster* : RNAi of $I_h$ : y[1] v[1]; P{y[+t7.7] v[+t1.8]=TRiP.JF03253}attP2 | Bloomington Drosophila Stock Center | BDSC:27574; FlyBase: FBst0029574 |
| *D. melanogaster* : RNAi of *Ork1* : P{KK107843}VIE-260B | Vienna *Drosophila* Resource Center | VDRC ID: 104883; FlyBase: FBst0476711 |
| *D. melanogaster* : RNAi of *Ork1* : y[1] v[1]; P{y[+t7.7] v[+t1.8]=TRiP.JF01926}attP2 | Bloomington Drosophila Stock Center | BDSC:25855; FlyBase: FBst0025885 |
| *D. melanogaster* : RNAi of *Shal* on the III chromosome | National Institute of Genetics Fly Stock Center | NIG Stock ID: 9262R-3 |
| *D. melanogaster* : RNAi of tipE on the III chromosome | National Institute of Genetics Fly Stock Center | NIG Stock ID: 1232R-3 |
| *D. melanogaster* : RNAi of *tipE* : y[1] v[1]; P{y[+t7.7] v[+t1.8]=TRiP.JF02148}attP2/TM3, Sb[1] | Bloomington Drosophila Stock Center | BDSC:26249; FlyBase: FBst0026249 |
| *D. melanogaster* : RNAi of *Atpα* : y[1] v[1]; P{y[+t7.7] v[+t1.8]=TRiP.JF02910}attP2 | Bloomington Drosophila Stock Center | BDSC: 28073; FlyBase: FBst0028073 |
| *D. melanogaster* : RNAi of *Atpα* : P{KK108782}VIE-260B | Vienna *Drosophila* Resource Center | VDRC ID: 100619; FlyBase: FBst0472492 |
| *D. melanogaster* : RNAi of *Calx* : y[1] v[1]; P{y[+t7.7] v[+t1.8]=TRiP.JF02937}attP2 | Bloomington Drosophila Stock Center | BDSC: 28306; FlyBase: FBst0028306 |
| *D. melanogaster* : RNAi of *Calx* : P{KK109144}VIE-260B | Vienna *Drosophila* Resource Center | VDRC ID: 104789; FlyBase: FBst0476622 |
| *D. melanogaster* : RNAi of *Ca-α1D* : y[1] v[1]; P{y[+t7.7] v[+t1.8]=TRiP.JF01848}attP2 | Bloomington Drosophila Stock Center | BDSC: 25830; FlyBase: FBst0025830 |
| *D. melanogaster* : RNAi of *Ca-α1D* : w[1118]; P{GD1737}v51490/CyO | Vienna *Drosophila* Resource Center | VDRC ID: 51490; FlyBase: FBst0469449 |
| *D. melanogaster:* RNAi of *Ca-α1T* : w[1118]; P{GD7754}v31963 | Vienna *Drosophila* Resource Center | VDRC ID: 31963; FlyBase: FBst0459316 |
| *D. melanogaster* : RNAi of *Ca-α1T* : y[1] v[1]; P{y[+t7.7] v[+t1.8]=TRiP.JF02150}attP2 | Bloomington Drosophila Stock Center | BDSC: 26251; FlyBase: FBst0026251 |
| *D. melanogaster* : RNAi of *eag* : y[1] v[1]; P{y[+t7.7] v[+t1.8]=TRiP.JF01471}attP2 | Bloomington Drosophila Stock Center | BDSC: 31678; FlyBase: FBst0031678 |
| *D. melanogaster* : RNAi of *eag* : P{KK107309}VIE-260B | Vienna *Drosophila* Resource Center | VDRC ID: 100260; FlyBase: FBst0472134 |
| *D. melanogaster* : RNAi of *Hk* : y[1] v[1]; P{y[+t7.7] v[+t1.8]=TRiP.JF02965}attP2/TM3, Sb[1] | Bloomington Drosophila Stock Center | BDSC: 28330; FlyBase: FBst0028330 |
| *D. melanogaster* : RNAi of *Hk* : P{KK109058}VIE-260B | Vienna *Drosophila* Resource Center | VDRC ID: 101402; FlyBase: FBst0473275 |
| *D. melanogaster* : RNAi of *inx2* : y[1] v[1]; P{y[+t7.7] v[+t1.8]=TRiP.JF02446}attP2 | Bloomington Drosophila Stock Center | BDSC: 29603; FlyBase: FBst0029306 |
| *D. melanogaster* : RNAi of *inx2* : P{KK111067}VIE-260B | Vienna *Drosophila* Resource Center | VDRC ID: 102194; FlyBase: FBst0474063 |
| *D. melanogaster* : RNAi of *Ir* : P{KK102249}VIE-260B | Vienna *Drosophila* Resource Center | VDRC ID: 107389; FlyBase: FBst0479211 |
| *D. melanogaster* : RNAi of *Irk2* : y[1] v[1]; P{y[+t7.7] v[+t1.8]=TRiP.JF01838}attP2 | Bloomington Drosophila Stock Center | BDSC: 25820;Flybase: FBst0025820 |

| | | |
|---|---|---|
| *D. melanogaster*: RNAi of *Irk2*: w[1118]; P{GD203}v4341 | Vienna *Drosophila* Resource Center | VDRC ID: 4341; FlyBase: FBst0465076 |
| *D. melanogaster*: RNAi of *KCNQ*: y[1] v[1]; P{y[+t7.7] v[+t1.8]=TRiP.JF02562}attP2 | Bloomington Drosophila Stock Center | BDSC: 27252; FlyBase: FBst0027252 |
| *D. melanogaster*: RNAi of *KCNQ*: P{KK109039}VIE-260B | Vienna *Drosophila* Resource Center | VDRC ID: 106655; FlyBase: FBst0478479 |
| *D. melanogaster*: RNAi of *Ncc69*: y[1] v[1]; P{y[+t7.7] v[+t1.8]=TRiP.JF03097}attP2 | Bloomington Drosophila Stock Center | BDSC: 28682; FlyBase: FBst0028682 |
| *D. melanogaster*: RNAi of *Ncc69*: P{KK108763}VIE-260B | Vienna *Drosophila* Resource Center | VDRC ID: 106499; FlyBase: FBst0478323 |
| *D. melanogaster*: RNAi of *Nckx30C*: y[1] v[1]; P{y[+t7.7] v[+t1.8]=TRiP.JF02574}attP2 | Bloomington Drosophila Stock Center | BDSC: 27246; FlyBase: FBst0027246 |
| *D. melanogaster*: RNAi of *nrv1*: w[1118] P{GD959}v46542 | Vienna *Drosophila* Resource Center | VDRC ID: 46542; FlyBase: Bst0466759 |
| *D. melanogaster*: RNAi of *nrv1*: P{KK100406}VIE-260B | Vienna *Drosophila* Resource Center | VDRC ID: 103702; FlyBase: FBst0475560 |
| *D. melanogaster*: RNAi of *nrv2*: y[1] v[1]; P{y[+t7.7] v[+t1.8]=TRiP.JF03081}attP2 | Bloomington Drosophila Stock Center | BDSC: 28666; FlyBase: FBst0028666 |
| *D. melanogaster*: RNAi of *nrv2*: w[1118]; P{GD960}v2660 | Vienna *Drosophila* Resource Center | VDRC ID: 2660; FlyBase: FBst0456497 |
| *D. melanogaster*: RNAi of *para*: w[1118]; P{GD3392}v6131 | Vienna *Drosophila* Resource Center | VDRC ID: 6131; FlyBase: FBst0470199 |
| *D. melanogaster*: RNAi of *para*: P{KK108534}VIE-260B | Vienna *Drosophila* Resource Center | VDRC ID: 104775; Flybase: FBst0476611 |
| *D. melanogaster*: RNAi of *picot*: y[1] v[1]; P{y[+t7.7] v[+t1.8]=TRiP.JF01940}attP2 | Bloomington Drosophila Stock Center | BDSC: 25920; FlyBase: FBst0025920 |
| *D. melanogaster*: RNAi of *picot*: P{KK106848}VIE-260B | Vienna *Drosophila* Resource Center | VDRC ID: 101082; FlyBase: FBst0472955 |
| *D. melanogaster*: RNAi of *ppk*: y[1] v[1]; P{y[+t7.7] v[+t1.8]=TRiP.JF03250}attP2 | Bloomington Drosophila Stock Center | BDSC: 29571; FlyBase: FBst0029571 |
| *D. melanogaster*: RNAi of *ppk*: P{KK104185}VIE-260B | Vienna *Drosophila* Resource Center | VDRC ID: 108683; FlyBase: FBst0480493 |
| *D. melanogaster*: RNAi of *ppk12*: y[1] v[1]; P{y[+t7.7] v[+t1.8]=TRiP.JF02027}attP2 | Bloomington Drosophila Stock Center | BDSC: 27092; FlyBase: FBst0027092 |
| *D. melanogaster*: RNAi of *ppk12*: P{KK101805}VIE-260B | Vienna *Drosophila* Resource Center | VDRC ID: 105131; FlyBase: FBst0476959 |
| *D. melanogaster*: RNAi of *ppk25*: y[1] v[1]; P{y[+t7.7] v[+t1.8]=TRiP.JF02434}attP2 | Bloomington Drosophila Stock Center | BDSC: 27088; FlyBase: FBst0027088 |
| *D. melanogaster*: RNAi of *ppk25*: P{KK109736}VIE-260B | Vienna *Drosophila* Resource Center | VDRC ID: 101808; FlyBase: FBst0473681 |
| *D. melanogaster*: RNAi of *ppk28*: y[1] v[1]; P{y[+t7.7] v[+t1.8]=TRiP.JF02153}attP2 | Bloomington Drosophila Stock Center | BDSC: 31878; FlyBase: FBst0031878 |
| *D. melanogaster*: RNAi of *ppk28*: P{KK106316}VIE-260B | Vienna *Drosophila* Resource Center | VDRC ID: 100946; FlyBase: FBst0472819 |
| *D. melanogaster*: RNAi of *sei*: y[1] v[1]; P{y[+t7.7] v[+t1.8]=TRiP.JF01474}attP2/TM3, Ser[1] | Bloomington Drosophila Stock Center | BDSC: 31681; FlyBase: FBst0031681 |
| *D. melanogaster*: RNAi of *sei*: P{KK105733}VIE-260B | Vienna *Drosophila* Resource Center | VDRC ID: 104698; FlyBase: FBst0476547 |
| *D. melanogaster*: RNAi of *Sh*: y[1] v[1]; P{y[+t7.7] v[+t1.8]=TRiP.JF01473}attP2/TM3, Ser[1] | Bloomington Drosophila Stock Center | BDSC: 31680; FlyBase: FBst0031680 |
| *D. melanogaster*: RNAi of *Sh*: P{KK109112}VIE-260B | Vienna *Drosophila* Resource Center | VDRC ID: 104474; FlyBase: FBst0476332 |
| *D. melanogaster*: RNAi of *Shal*: y[1] v[1]; P{y[+t7.7] v[+t1.8]=TRiP.JF02154}attP2 | Bloomington Drosophila Stock Center | BDSC: 31879; FlyBase: FBst0031879 |
| *D. melanogaster*: RNAi of *Shaw*: y[1] v[1]; P{y[+t7.7] v[+t1.8]=TRiP.JF02982}attP2 | Bloomington Drosophila Stock Center | BDSC: 28346; FlyBase: FBst0028346 |
| *D. melanogaster*: RNAi of *Shaw*: P{KK108371}VIE-260B | Vienna *Drosophila* Resource Center | VDRC ID: 110589; FlyBase: FBst0482154 |
| *D. melanogaster*: RNAi of *SK*: y[1] v[1]; P{y[+t7.7] v[+t1.8]=TRiP.JF02571}attP2 | Bloomington Drosophila Stock Center | BDSC: 27238; FlyBase: FBst0027238 |

| | | |
|---|---|---|
| *D. melanogaster* : RNAi of *SK* : P{KK107699}VIE-260B | Vienna *Drosophila* Resource Center | VDRC ID: 103985; FlyBase: FBst0475843 |
| *D. melanogaster* : RNAi of *SLO2* : y[1] v[1]; P{y[+t7.7] v[+t1.8]=TRiP.JF03426}attP2 | Bloomington Drosophila Stock Center | BDSC: 32034; FlyBase: FBst0032034 |
| *D. melanogaster* : RNAi of *stj* : y[1] v[1]; P{y[+t7.7] v[+t1.8]=TRiP.JF01825}attP2 | Bloomington Drosophila Stock Center | BDSC: 25807;FlyBase: FBst0025807 |
| *D. melanogaster* : RNAi of *stj* : P{KK101267}VIE-260B | Vienna *Drosophila* Resource Center | VDRC ID: 108569; FlyBase: FBst0480379 |
| *D. melanogaster* : RNAi of *Teh2* : w[1118]; P{GD3839}v9037 | Vienna *Drosophila* Resource Center | VDRC ID: 9037; FlyBase: FBst0471346 |
| *D. melanogaster* : RNAi of *Teh2* : P{KK112449}VIE-260B | Vienna *Drosophila* Resource Center | VDRC ID: 104951; FlyBase: FBst0476779 |
| *D. melanogaster* : RNAi of *Teh4* : w[1118]; P{GD3578}v11621/CyO | Vienna *Drosophila* Resource Center | VDRC ID: 11621; FlyBase: FBst0450303 |
| *D. melanogaster* : RNAi of *Teh4* : P{KK110985}VIE-260B | Vienna *Drosophila* Resource Center | VDRC ID: 102161; FlyBase: FBst0474030 |
| *D. melanogaster* : RNAi of *trp* : y[1] v[1]; P{y[+t7.7] v[+t1.8]=TRiP.JF01441}attP2 | Bloomington Drosophila Stock Center | BDSC: 31649; FlyBase: FBst0031649 |
| *D. melanogaster* : RNAi of *trp* : w[1118]; P{GD372}v1366 | Vienna *Drosophila* Resource Center | VDRC ID: 1366; FlyBase: FBst0451102 |
| *D. melanogaster* : UAS-*pdf* on the 2$^{nd}$ Chromosome | Renn et al., 1999 | Not available |
| **Software** | | |
| ImageJ | Schneider et al., 2012 | https://imagej.nih.gov/ij/ |
| Infostat | Di Rienzo et al., 2013 | https://www.infostat.com.ar/ |
| ClockLab | Actimetrics | https://www.actimetrics.com/products/clocklab/ |
| Rethomics | Geissmann et al., 2019 | https://rethomics.github.io/ |
| R | R Core Team, 2014. | https://www.r-project.org/ |
| Micro Manager | Edelstein et al., 2014 | https://micro-manager.org/wiki/Download_Micro-Manager_Latest_Release |
| pClamp 9 | Molecular Devices | https://moleculardevices.app.box.com/s/d93nukl3chbo206t33cw5fpabsph6wh4 |
| Clampfit 10 | Molecular Devices | https://moleculardevices.app.box.com/s/l8h8odzbdikalbje1iwj85x88004f588 |
| Origin 8 | OriginLab | https://www.originlab.com/ |
| Graphpad | Prism8 | https://www.graphpad.com/ |

**S2 Table: Reagents and resources used for this work.**
